# Neural basis of concurrent deliberation toward a choice and confidence judgment

**DOI:** 10.1101/2024.08.06.606833

**Authors:** Miguel Vivar-Lazo, Christopher R. Fetsch

**Affiliations:** Zanvyl Krieger Mind/Brain Institute, Johns Hopkins University, Baltimore, MD, USA; Solomon H. Snyder Department of Neuroscience, Johns Hopkins University, Baltimore, MD, USA

## Abstract

Decision confidence plays a key role in flexible behavior and (meta)cognition, but its underlying neural mechanisms remain elusive. To uncover the latent dynamics of confidence formation at the level of single neurons and population activity, we trained nonhuman primates to report choice and confidence with a single eye movement on every trial. Monkey behavior was well fit by a bounded accumulator model instantiating parallel processing of evidence, rejecting a serial model in which the choice is resolved first followed by post-decision accumulation for confidence. Neurons in area LIP reflected concurrent accumulation, showing covariation of choice and confidence signals across the population, and within-trial dynamics consistent with parallel updating at near-zero time lag. The results demonstrate that the primate brain can process a single stream of evidence in service of two computational goals simultaneously—a categorical decision and associated level of confidence—and illuminate a candidate neural substrate for this capacity.

## INTRODUCTION

Evolution has endowed humans and some animals with the ability to assess the quality of their own decisions. This typically manifests as a degree of confidence, defined as an internal estimate of the probability of being correct. Confidence facilitates learning in the absence of explicit feedback (Daniel & Pollmann, 2012; Guggenmos et al., 2016) and guides decisions that are part of a sequence or hierarchy (Sarafyazd & Jazayeri, 2019; Purcell & Kiani, 2016; van den Berg et al., 2016b; Zylberberg, 2021). When feedback does occur, confidence informs whether the outcome is surprising (i.e., a high confidence error), driving a change in learning rate (Rescorla & Wagner, 1972; Sutton & Barto, 1992). Indeed it can be shown that the optimal weights for converting sensory neuron activity to decision evidence can only be obtained with a learning rule that is proportional to confidence (Drugowitsch et al., 2019). To pursue a mechanistic understanding of these downstream effects, it is critical to establish how, where, and when neural representations of confidence emerge in the brain. In particular, the timing of confidence computations (Xue & Rahnev, 2023) within and across brain regions constrains the flow of information underlying its effects on learning and sequential decision making, and is an important component of more general theories of metacognition (Fleming, 2024).

In the context of dynamic (i.e., evidence accumulation) models, there are three main possibilities for the relative timing of choice and confidence formation. *Serial* models posit an initial phase of accumulation bearing only on the choice, followed by a secondary process that governs confidence (Pleskac & Busemeyer, 2010; Moran et al., 2015; Herregods et al., 2024). Although the readout of confidence from the second process is conditioned on which choice bound was reached, it is otherwise agnostic to the primary accumulation epoch, and there is no mechanism for reading out a provisional degree of confidence while the decision is still being formed. In contrast, *parallel* models propose a simultaneous initiation and temporally overlapping process for choice and confidence, with an explicit mapping between the state of the accumulator(s) at each point in time and the probability of being correct if the decision were to terminate there (Kiani et al., 2014a; van den Berg et al., 2016a; Khalvati et al., 2021). Lastly, within parallel models we can define a subcategory (*hybrid*) as having an initial parallel phase followed by a period of post-decision accumulation that only affects confidence (Desender et al., 2021a; Maniscalco et al., 2021; Hellmann et al., 2023; Le Denmat et al., 2024). Note that the serial/parallel question is orthogonal to whether confidence and choice derive from distinct transformations of the evidence (Maniscalco et al., 2021; Hellman et al., 2023; Balsdon et al., 2020, Balsdon & Philiastides 2024); we return to this important issue in the Discussion.

Recent behavioral studies in humans (Dotan et al., 2018; Balsdon et al., 2020; Li et al., 2023) offer compelling evidence for parallel computation of confidence, further supported by electroencephalography (Gherman & Philiastides, 2015; Balsdon et al., 2021; Balsdon & Philiastides 2024; Dou et al., 2024; Goueytes et al., 2024) and transcranial magnetic stimulation (Xue et al., 2023). Whether nonhuman animals possess this ability, and how it might be implemented at the level of neuronal populations, remains unclear. We addressed this gap by training monkeys to report choice and confidence simultaneously in a reaction-time (RT) paradigm (‘peri-decision wagering’, peri-dw), building on previous work in human participants (Kiani et al., 2014a; van den Berg et al., 2016a). Our goal was not to question the importance of post-decisional processing in general, but to ask if a serial process is essential for a monkey’s deliberation about both aspects of the decision or if they can occur in parallel. Notably, unlike ‘opt-out’ or ‘uncertain-response’ paradigms (Kiani et al., 2009; Smith et al., 2012; Komura et al., 2013; Fetsch et al., 2014a; Li et al., 2023), the peri-dw task affords direct measurement of choice, RT, and confidence and their relationship to neural activity on trial-by-trial basis (see also Boundy-Singer et al., 2024).

While monkeys performed the task, we recorded population spiking activity in the ventral portion of the lateral intraparietal area, LIPv (Shadlen & Kiani, 2013). Previous work has shown that LIPv (hereafter LIP) represents a decision variable (DV) that predicts choice and RT (Roitman & Shadlen, 2002), as well as confidence in an opt-out task (Kiani & Shadlen, 2009). We found that behavior in the peri-dw task is best explained by concurrent evaluation of evidence for choice and confidence, and that neural activity in LIP reflects the requisite parallel process. The findings support a role for posterior parietal cortex in behaviors guided by an online estimate of confidence, and more broadly favor an architecture for visual metacognition that is fundamentally parallel.

## RESULTS

We recorded 407 neurons in area LIP in the right hemisphere of two rhesus monkeys (*Macaca mulatta*; 207 in monkey H, 200 in monkey G) while they performed the peri-dw task (Fig. 1a). Each saccade target corresponds to a motion direction judgment (left or right) and a wager (high or low) on the correctness of that judgment. Although behaviorally the task amounts to a single choice among four options, we refer to the left-right component as ‘choice’ and the high-low component as ‘wager’ or ‘bet’, both for simplicity and because the results support this interpretation. Monkeys were rewarded or penalized based on the conjunction of accuracy and wager (Fig. 1b): a larger drop of juice for high vs. low bets when correct, and a time penalty for high bets when incorrect (no penalty for a low-bet error).

**Figure 1.**
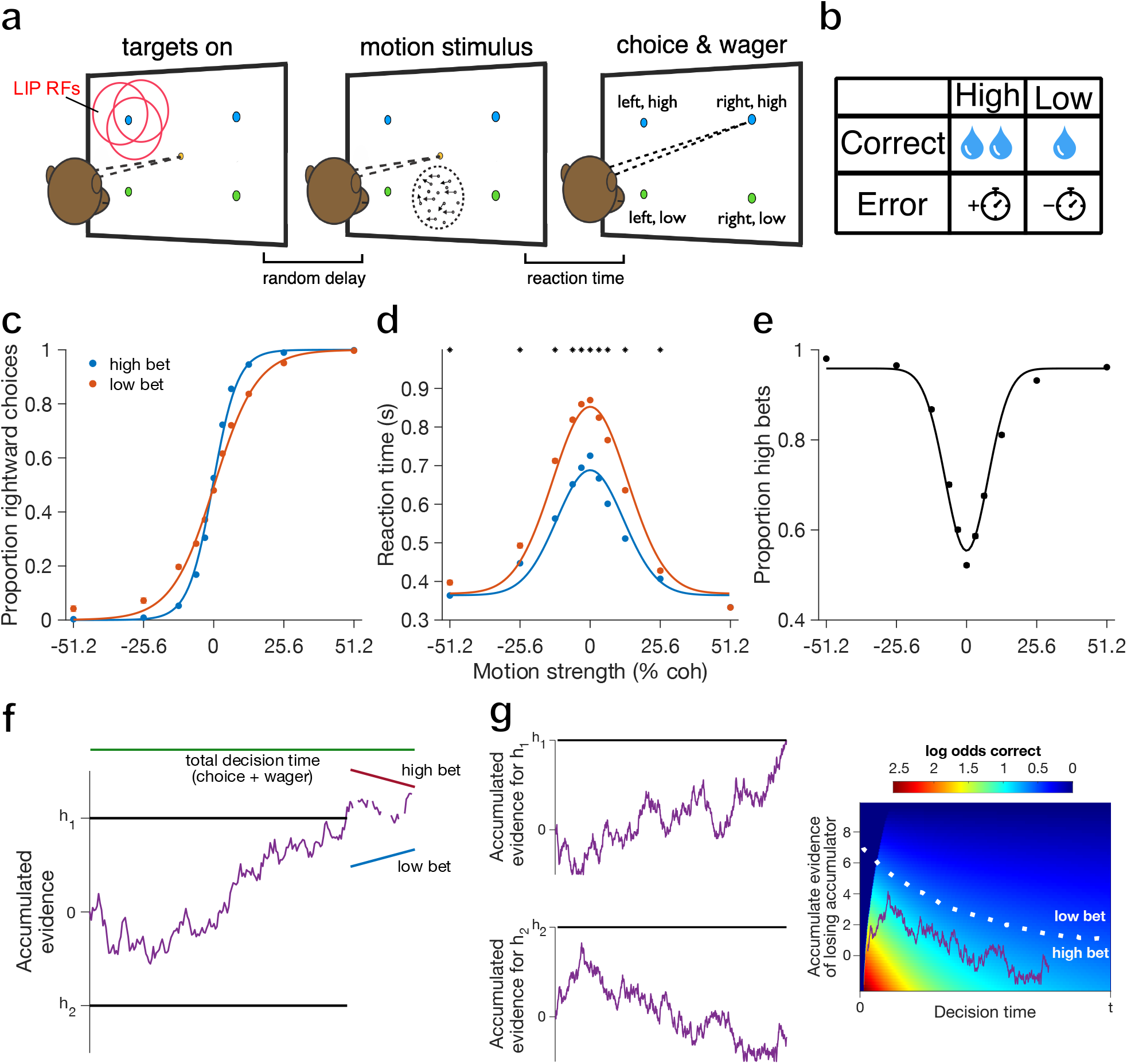
Task sequence, behavioral performance and model schematics. **a**, After the monkey acquires fixation, four targets are presented, followed by a random-dot motion stimulus. At any time after motion onset the monkey can make a saccade to one of the targets to indicate its choice and wager. **b**, Table showing the possible outcomes for each trial: if correct, a high bet yields a larger juice reward compared to a low bet, but if incorrect, a high bet incurs a 2-3 s time penalty (added to the next trial’s pre-stimulus fixation period). Low-bet errors were not penalized. **c**-**e**, Behavioral results pooled across two monkeys (N = 372 sessions, 401,530 trials, including sessions without neural recording). Each ordinate variable is plotted as a function of signed motion strength (%coh, negative=leftward, positive=rightward). Choice and RT functions are shown conditioned on the wager (low=red, high=blue). Error bars (SE) are smaller than the data points. Smooth curves show logistic regression (choice) and Gaussian (RT, wager) fits. **f**, The serial model begins with a single accumulator with symmetric bounds (standard 1D drift-diffusion model). Arriving at one of the bounds terminates the primary decision (h_1_ vs. h_2_ = left vs. right in our task) and initiates a secondary process that accumulates evidence toward a ‘high’ or ‘low’ bound governing the wager. **g**, The parallel model comprises two concurrent accumulators (left) that are partially anticorrelated. The first bound to be crossed (the ‘winner’) dictates the choice and decision time, whereas the losing accumulator dictates confidence by way of a mapping (right) between accumulated evidence and the log odds that the choice made was the correct one (color scale).

As in previous studies (e.g., Roitman & Shadlen, 2002), monkeys showed greater accuracy (Fig. 1c) and faster RTs (Fig. 1d) when the motion was strong compared to weak (coh near 0%). Motion strength also influenced wagering behavior in a sensible manner, namely the probability of betting high increased with greater motion strength in either direction (Fig. 1e). Importantly, the behavior shows that the low-bet option did not correspond to opting out or failing to engage in the motion decision: accuracy remained high on low-bet trials, and choice and RT still varied systematically with motion strength in a manner consistent with a deliberative process (see Model Fitting below).

As expected for a behavioral assay of confidence, the monkeys’ sensitivity was greater when betting high versus low (Fig. 1c, red vs. blue; T = 34.41, p = 1.65×10^−259^, se = 0.2735, logistic regression). This was true even when controlling for variability in motion energy within each coherence level, by leveraging multiple repeats of the same random seed (Bondy et al., 2018; Supplementary Fig. 1a; monkey H: p = 1.3376×10^−04^, T = 3.8194; monkey G: p = 3.4828×10^−07^, T = 5.0953; see Methods). Both monkeys also showed faster RTs when betting high versus low, for all but the largest motion strengths (Fig. 1d, red vs. blue; asterisks indicate p < 0.0045 by t-test, Bonferroni corrected). This improved sensitivity and faster RTs for high-bet choices was evident in the large majority of individual sessions, as assessed with logistic regression (Extended Data Fig. 1a; monkey H: p = 2.8388×10^−20^, T = 11.63, CI = [7.9377 Inf]; monkey G: p=9.4419×10^−38^, T=18.74, CI = [8.4779 Inf]; one-tailed t-test) and Gaussian fitting (Extended Data Fig. 1b; monkey H: p = 5.0392×10^−08^, T = −5.76, CI = [−Inf −0.053]; monkey G: p = 1.4136×10^−25^, T = −13.26, CI = [ −Inf −0.241]; one-tailed t-test). Lastly, we examined wagering behavior as a function of RT quantile, separately for each individual motion strength (Extended Data Fig. 2a-b). For most motion strengths, the monkeys bet high less often for longer RTs (Extended Data Fig. 2b; p < 0.0085 for both monkeys for every coherence except 51.2%, Cochran-Armitage test with Bonferroni correction). This pattern is strikingly similar to that observed in humans on a similar task (Kiani et al., 2014a), where the results suggested that confidence depends on both evidence strength and elapsed time. As in that previous study, the pattern remained significant when controlling for variability in motion energy across trials of a given coherence (Extended Fig. 2d; monkey H: p = 3.2351×10^−04^, F = 5.24; monkey G: p=0.0056, F = 3.65; interaction term between motion energy and RT quintile using ANCOVA). An inverse relationship between RT and confidence is a classic psychophysical result (Henmon, 1911; Kellogg, 1931; Audley, 1960), replicated in more recent human work (e.g., Kiani et al., 2014a; Desender et al., 2021a; Dou et al., 2024). Observing it in monkeys, for the first time to our knowledge, supports the peri-dw assay as a valid measure of confidence, and it is consistent with a family of accumulator models as described below.

### Model-free analyses reveal temporal overlap in choice and confidence computations

Although the choice and wager were indicated with a single eye movement, this does not necessitate simultaneity in the processing of evidence. Different temporal windows of the stimulus could covertly be used to support the two elements of the decision, which would then only be reported when both were resolved. To test whether monkeys use a consistent serial strategy (resolving choice first and then confidence, or vice versa) we calculated the influence of stimulus fluctuations on choice and confidence as a function of time (psychophysical kernels; Kiani et al., 2008; Nienborg & Cumming, 2009; Zylberberg et al., 2012a). Briefly, we quantified the motion energy for each trial and video frame by convolving the random-dot pattern with two pairs of spatiotemporal filters aligned to leftward and rightward motion (Adelson & Bergen, 1985). We then partitioned trials by outcome and plotted the average relative motion energy (residuals) for each outcome as a function of time.

Psychophysical kernels for choice are plotted in Fig. 2a-b. Rightward choices were preceded by more rightward motion energy throughout most of the trial (red line), and the same was true for leftward choices and leftward motion (blue). The kernels for right and left choice began to separate about 100 ms after motion onset and remained so until ~100 ms before saccade initiation. This clear separation suggests that the monkeys used essentially the entire stimulus epoch, on average, to decide motion direction. For confidence, we calculated the kernels by taking the difference between motion energy time series for high and low bets associated with a specific choice (van den Berg, 2016a). For high minus low wager on rightward choice trials, motion energy values were above zero, indicating an excess of rightward motion on high-bet choices compared to low-bet choices (Fig. 2c-d, green). Similarly, for left choice trials the difference in motion energy was below zero, indicating more leftward motion on high vs. low bets (Fig. 2c-d, purple). This analysis shows that both early and late motion evidence is leveraged to inform confidence, in both monkeys. Comparison of the traces in Figure 2a-b vs. 2c-d might suggest that the utilization of the stimulus for confidence does not identically overlap with choice, especially for monkey H. However, the substantial overlap does appear to rule out a consistent temporal segregation, such as an obligatory post-decision mechanism for confidence.

**Figure 2.**
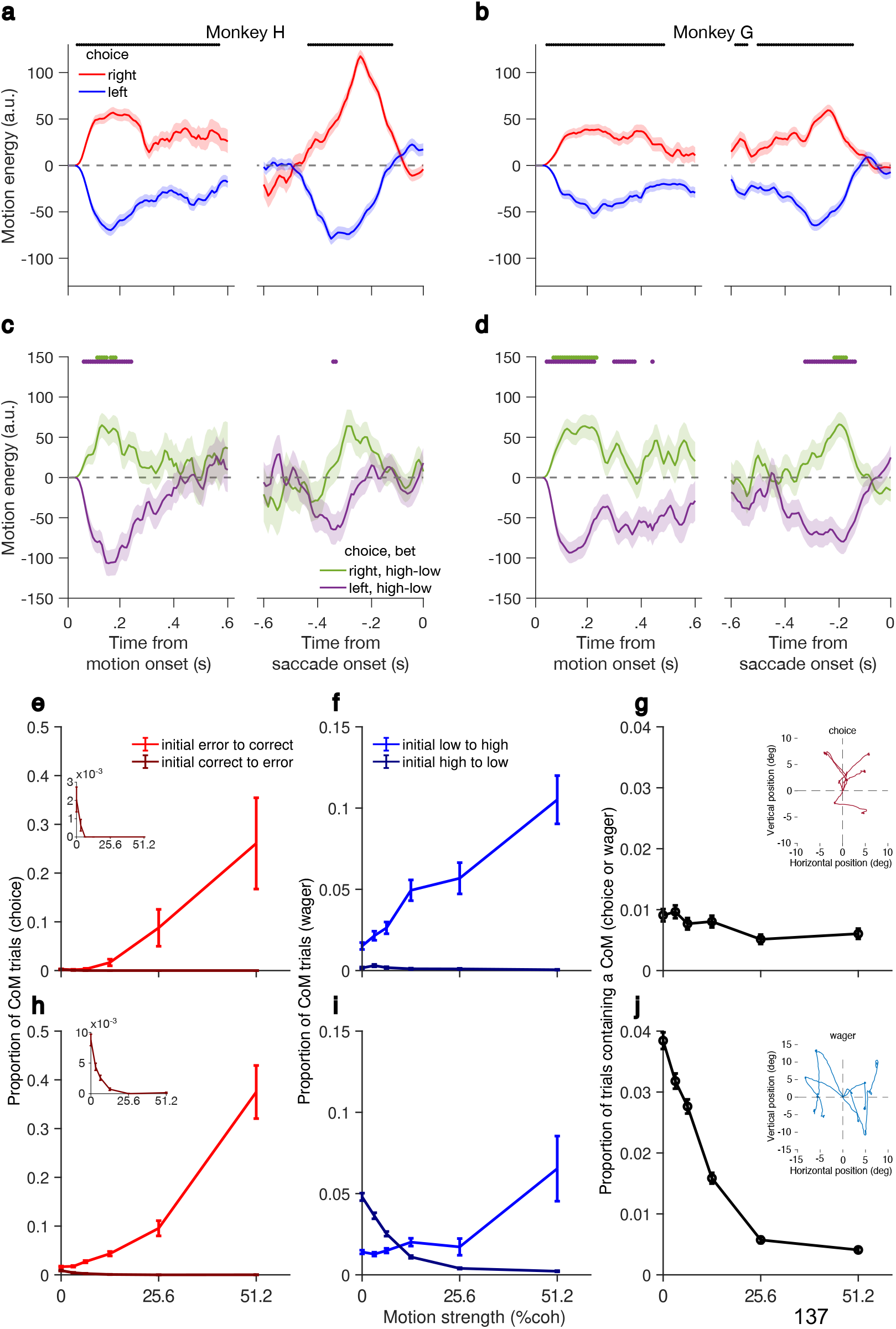
Psychophysical kernels and changes of mind for choice and wager. **a**,**b**, Motion energy profiles conditioned on right and left choices (red and blue, respectively), aligned to motion onset and saccade onset. Shaded regions indicate SEM. Black line at top indicates when right and left traces were significantly different from each other (p<0.05, t-test with Šidák Correction for 140 frames). **c**,**d**, Confidence kernels computed as the difference in motion energy between right-high and right-low choices (green), and the difference between left-high and left-low choices (purple), aligned to the same events as **a**,**b**. Colored lines at top of graph indicate when the corresponding traces were significantly different from zero (p<0.05, t-test with Šidák correction). **e**,**h**, Proportion of change-of-mind (CoM) trials that start with an initial error (light red) or correct (dark red, y-axis expanded in insets) and end as correct or error, respectively, as a function of motion strength. In e-j, top row shows data from monkey H and bottom row is monkey G. **f**,**i**, Proportion of trials that start with an initial low wager (light blue) or high wager (dark blue) and end as high or low wager, respectively, as a function of motion strength. **g**,**j**, Proportion of trials that are CoMs (of choice, wager, or both) as a function of motion strength. Insets show example eye trajectories from choice (top) and wager (bottom) CoM trials.

When decisions are reported with an arm movement, subjects occasionally alter their reach trajectory in a manner that suggests a ‘change of mind’ (CoM) based on continued processing of evidence after movement initiation (Resulaj et al., 2009; van den Berg et al., 2016a). Although saccadic choices, being fast and ballistic, are often assumed to be incompatible with CoMs (but see McPeek et al., 2000; Caspi et al., 2004), we identified a small subset of trials with multiple saccades in quick succession that showed certain characteristic features (Fig. 2e-j). These putative CoMs were more frequent on difficult vs. easy trials (Fig. 2g & 2j; monkey H: p = 0.007, T = −3.70; monkey G: p = 3.1983×10^−07^, T = −31.15, Cochran-Armitage test), and changes from incorrect to correct were more likely when motion strength was high (Fig. 2e & 2h red line; monkey H: p = 5.7576×10^−05^, T = 10.86; monkey G: p = 1.5973×10^−05^, T = 14.13). Correct-to-error CoMs, occurring sparingly, were more likely when motion strength was low (Fig. 2e & 2h, dark red line; monkey H: p = 0.001, T = −5.88; monkey G: p = 6.9991×10^−06^, T = −16.72). We also observed changes from low to high confidence, which for both monkeys were more frequent with greater motion strength (Fig. 2f & 2i blue line; monkey H: p = 4.9597×10^−05^, T = 11.20; monkey G: p = 0.01, T = 3.38), as shown previously in humans (van den Berg et al., 2016a). The presence of CoMs and changes of confidence, sometimes both on the same trial, implies that both aspects of the decision were subject to revision at the time of the initial saccade. This is inconsistent with a strictly serial process, though it also reveals a brief window for post-decision processing even for saccadic choices (McPeek et al., 2000; Caspi et al., 2004).

### Model fitting supports parallel deliberation for choice and confidence

Previous studies have typically adopted a particular temporal framework rather than comparing across model classes (but see Hellmann et al., 2023; Shekhar & Rahnev, 2024). Here we provide a thorough comparison of serial, parallel, and hybrid models fitted to the same data from the peri-dw task. The parallel and hybrid models consist of two accumulators integrating evidence for the two motion directions, differing only in when accumulated evidence is mapped to confidence. To explain the wager, the confidence mapping is binarized by a single free parameter: a criterion on log odds correct associated with a high vs. a low bet (Fig. 1g, right). The hybrid model adds an additional free parameter for the duration of post-decision accumulation. For the serial model we made the simplifying assumption (as in previous work, refs. Pleskac & Buseyemyer, 2010; Moran et al., 2015; Herregods et al., 2024) that the two accumulators were perfectly anticorrelated, equivalent to a one-dimensional drift diffusion model (DDM). After one of the two bounds is reached, evidence continues to accumulate towards a second set of bounds dictating the wager (Fig. 1f). The observed RT is the sum of the time taken to reach both bounds, plus non-decision time.

The smooth curves in Figure 3 and Extended Data Figure 3 are fits to the serial, parallel, and hybrid models for both monkeys. All models perform quite well at describing choice, RT, and confidence as a function of motion strength when pooled across correct/incorrect and high/low wager trials (Extended Data Fig. 3), a testament to the explanatory power of the bounded accumulation framework. Interestingly, all three models also qualitatively capture the greater choice sensitivity and faster RTs for high vs. low wager (Fig. 3, first and second column). This comparison illustrates the difficulty of disambiguating the mechanism(s) underlying choice and confidence using behavior alone. Indeed, quantitative model comparison yielded mixed results for the two monkeys: hybrid and parallel models were favored over the serial model for monkey G (Bayesian Information Criterion: hybrid = 1.1610×10^6^, parallel = 1.1614×10^6^, serial = 1.1644×10^6^), but the opposite was true for monkey H (BIC: serial = 7.7993×10^5^, parallel = 7.8179×10^5^, hybrid = 7.8279×10^5^).

**Figure 3.**
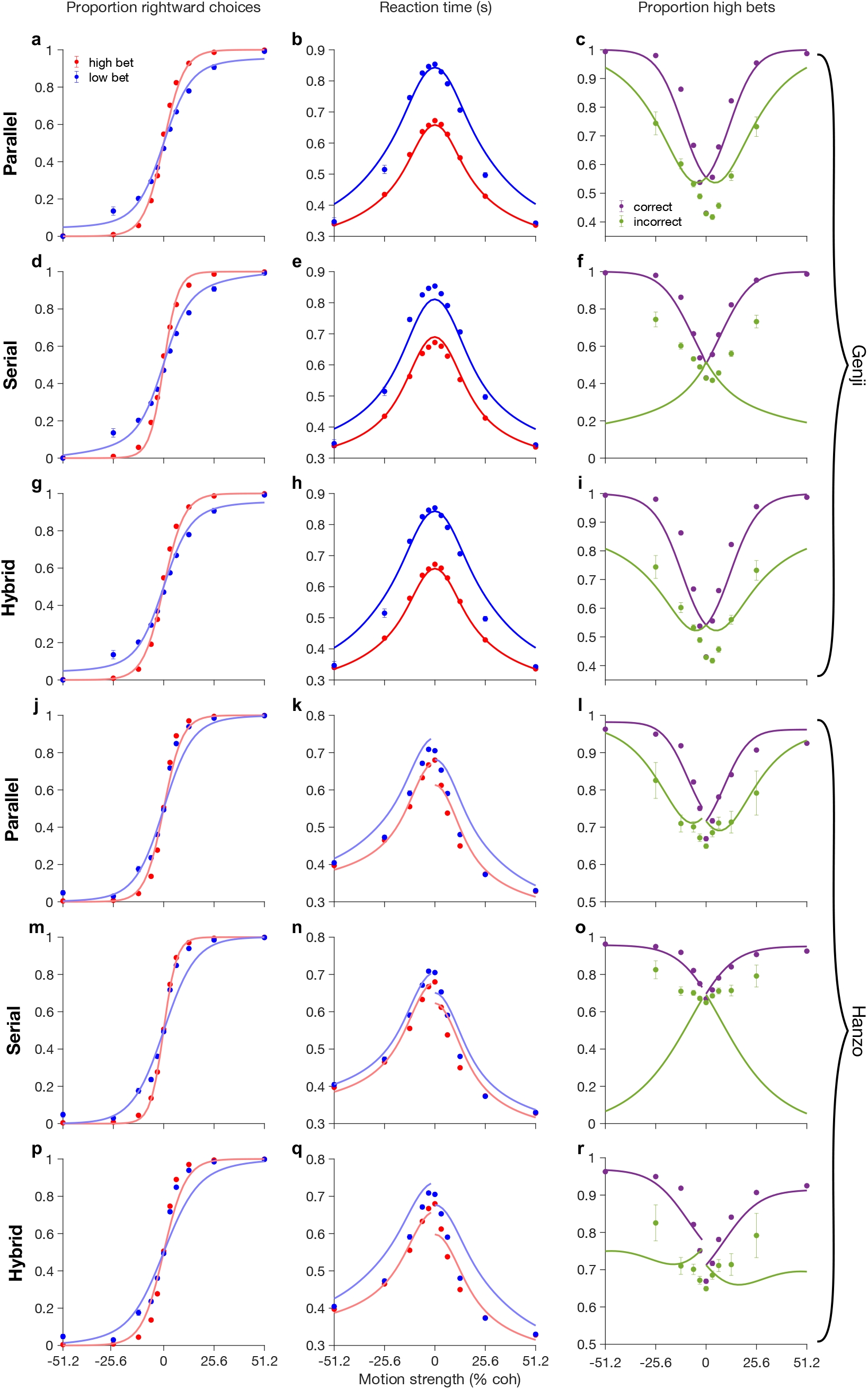
Comparison of parallel, serial, and hybrid model fits. Left column shows proportion of rightward choices as a function of motion strength (%coh), conditioned on high and low bet trials (red and blue, respectively). Middle column shows mean reaction time in the same format. Right column shows proportion of high bets as a function of motion strength (% coh), conditioned on correct vs. error trials (magenta and green, respectively). Panels **a**-**i** correspond to the results from monkey G. **a**-**c**, Fits to the parallel model. **d**-**f**, Fits to the serial model. **g**-**i**, Fits to the hybrid model. The solid dots are empirical data points (+/-S.E) and solid lines are model fits. In many cases the error bars are smaller than the data points. **j**-**r**, Same as **a**-**i** but for monkey H.

Critically, however, the serial model fails in one key aspect: the pattern of wagering behavior conditioned on accuracy (Fig. 3, right column). It is commonly observed that confidence ratings increase as a function of evidence strength on correct trials, but *decrease* with evidence strength on incorrect trials. This characteristic ‘X’ pattern (or ‘folded-X’, if stimulus strength is unsigned) has been proposed as a statistical signature of confidence in both behavior and brain activity (Kepecs et al., 2008; Rolls et al., 2010; Komura et al., 2013; Sanders et al., 2016; Bang et al., 2018), yet it is not universal. Other studies (Kiani et al., 2014a; van den Berg et al., 2016a; Rausch et al., 2018) report that confidence increases with evidence strength even for errors, and this is what we observed as well (Fig. 3, right column). It is becoming increasingly clear that these conflicting findings can, in many cases, be explained by a temporal dissociation (Kiani et al., 2014a; Fetsch et al., 2014b; van den Berg et al., 2016a; Desender et al., 2021a; Khalvati et al., 2021): resolving the choice first followed by confidence later allows for revision of the confidence judgment upon further deliberation. Incorrect choices when the stimulus was strong are more likely to undergo such revision, hence confidence decreases (on average) with evidence strength on error trials. Reducing or eliminating the delay between the choice and confidence report tends to flatten or reverse the X pattern (Kiani et al., 2014a; van den Berg, 2016a, Desender et al., 2021a). Because the serial and hybrid models impose such a delay (implicitly, in our task), they cannot reproduce the qualitative trend in error-trial confidence we observed empirically (Fig. 3, right column, green data points)—unless the post-decision epoch is very brief, as it was in the best-fitting hybrid model for monkey G (60 ms; Fig. 3i). This qualitative miss is not reflected in the above BIC results because the model likelihoods were calculated using the unconditioned wager data, meaning the split between correct and error trials (Fig. 3, right column, green vs. purple curves) is a prediction, not a fit. Quantifying the quality of this prediction using the error-trial wagers establishes the parallel and serial model as the most and least supported, respectively, for both monkeys (negative log-LL: parallel = 8.4320×10^3^, hybrid = 8.4370×10^3^, serial 8.6572×10^3^ for monkey H; parallel = 1.5636×10^4^, hybrid = 1.5699×10^4^, serial = 1.5993×10^4^ for monkey G).

In summary, although each of the model variants is flexible enough to capture most behavioral trends, a holistic model comparison favors a process where evidence is accumulated in parallel for constructing a decision and associated level of confidence. We next examine whether decision-related activity in parietal cortex is consistent with such a process.

### LIP neurons show signatures of concurrent accumulation

Putative DV representations can be found in many subcortical and cortical areas, characterized by a ‘ramping’ pattern of neural activity (or decoded proxy thereof) that scales with evidence strength (and/or response time) and often converges upon decision termination (Shadlen & Kiani, 2013; Yartsev et al., 2018; O’Connell & Kelly, 2021; Bondy et al., 2024). Although this pattern does not uniquely identify a process of evidence accumulation, a large body of work supports the assertion that macaque LIP neurons reflect such a process during the random-dot motion task (Shadlen & Kiani, 2013; Steinemann et al., 2024). We reasoned that, if choice and confidence were resolved concurrently during motion viewing (parallel model), the ramping activity should begin to predict both dimensions of the eventual saccade at the same time, classically around 200 ms after motion onset (Roitman & Shadlen, 2002). Alternatively, if choice were deliberated first followed by confidence (serial model), this temporal separation should be evident in the divergence point of neural activity traces conditioned on the four outcomes.

These traces are shown in Figure 4a for four example neurons. The highest firing rate corresponds to choices made into the receptive field (RF) of the neuron, which was almost always in the left (contralateral) hemifield but was equally likely to overlap the high or low wager target. The relative ordering of the remaining three traces differs across neurons, possibly due to idiosyncratic RF properties or nonspatial decision signals. The key observation is that the activity preceding saccades to the preferred wager target (low or high) diverges from the activity for the other wager target (high or low) at about the same time as it diverges from the traces for ipsilateral choice (right-low and right-high). This pattern is present in each example neuron as well as in the population averages (Fig. 4b-c). There is no evidence that ramping activity consistently predicts the left-right choice sooner than the high-low one (or vice versa), as expected under a serial model. Instead, to the extent the activity reflects accumulation of evidence favoring the target in the RF (see below and Discussion), the results support a model in which such accumulation underlies concurrent deliberation toward a choice and confidence judgment.

**Figure 4.**
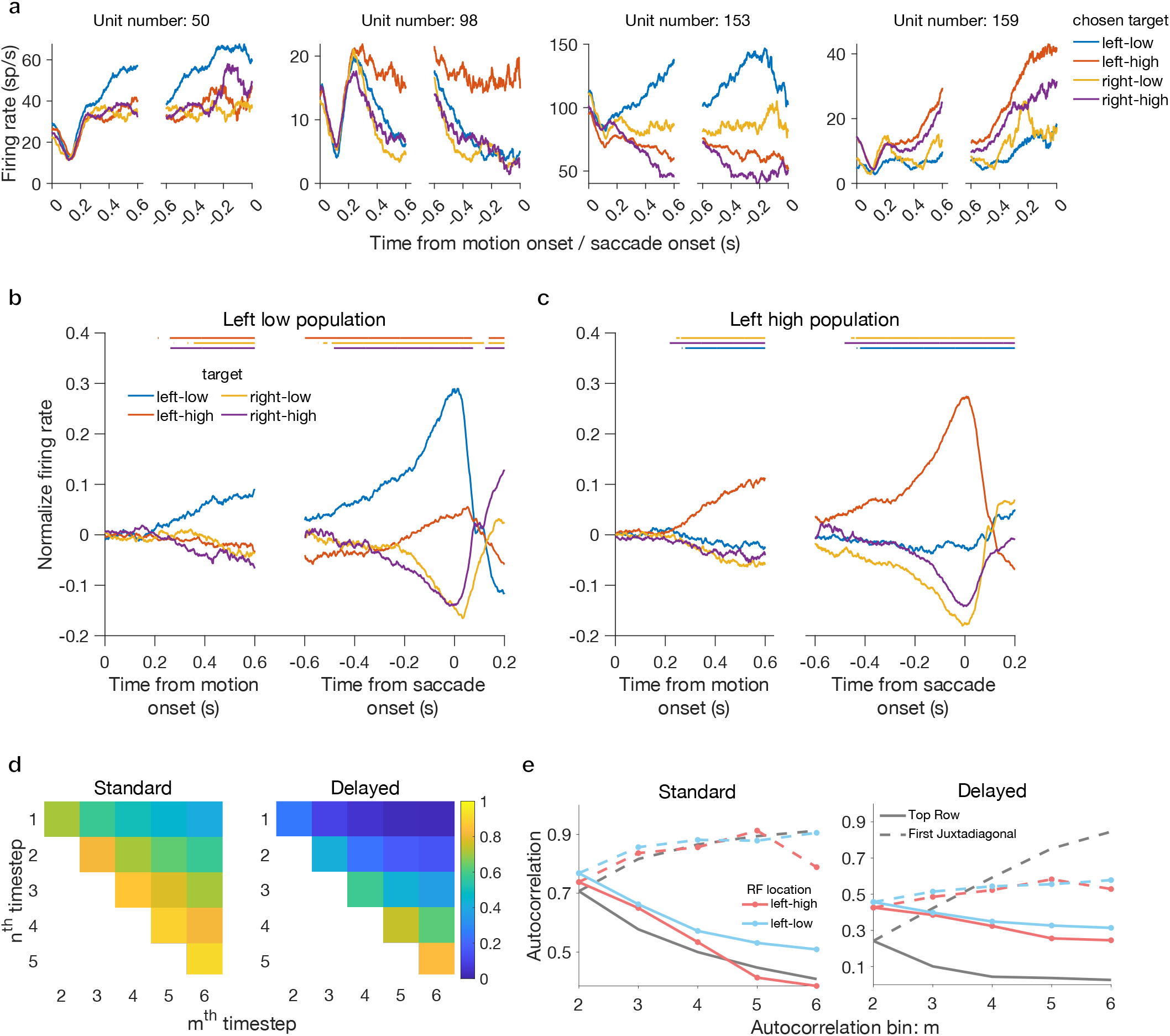
Temporal properties of neural activity in LIP during the peri-dw task. **a**, Firing rate of example units split by choice/wager outcome, aligned to motion onset and saccade onset. **b**, Population average firing rate (normalized) for neurons with an RF overlapping the left-low target. Colored bars at top indicate when the corresponding trace is significantly below the trace for choices into the RF. **c**, Same as **b** but for left-high neurons. Only low-coherence trials (0, +/-3.2%, +/-6.4%) are included in **a**-**c. d**, Theoretical autocorrelation matrix of a standard ideal accumulation process (left) and a delayed accumulator (right; see Methods). **e**, Projection of theoretical autocorrelations for top row (gray dotted) and first juxtadiagonal (gray solid) along with the corresponding empirical data after fitting the phi parameter (Methods). Blue and Red traces represent the two populations shown in **b** and **c**, respectively, pooling the data from both monkeys. Left and right panels show standard and delayed accumulators, respectively. The empirical traces (blue and red) are not identical for Standard and Delayed because the phi parameter is fit independently (see Methods).

To dig deeper into the nature of the observed ramping signals, we tested for statistical signatures of a bounded accumulation process (Churchland et al., 2011; de Lafuente et al., 2015; Shushruth et al., 2018; Steinemann et al., 2024): (a) increasing variance of the underlying rate (variance of conditional expectation, VarCE) followed by a collapse at decision time, and (b) a characteristic autocorrelation pattern in this latent signal (correlation of conditional expectation, CorCE; see Methods). The results supported both sets of predictions. After 200 ms following motion onset, VarCE shows a roughly linear increase for at least the next 400 ms (Extended Data Fig. 4, left; left-high neurons, monkey H: p < 10^−7^, monkey G: p < 10^−13^; left-low neurons, monkey H: p < 10^−19^, monkey G: p = 0.0014; linear regression), then decreases near saccade initiation (Extended Data Fig. 4, right; left-high neurons, monkey H: p = 0.015, monkey G: p = 0.0016; left-low neurons, monkey H: p < 10^−4^, monkey G: p < 10^−7^). For CorCE, the results from both monkeys were reasonably well matched to the predictions, namely an increase in the correlation between neighboring time bins as time elapses, and a decrease in correlation between bins as the separation between them increases (Fig. 4d & e, left; monkey G: R^2^ = 0.74 and 0.79 for left-high and left-low neurons, respectively; monkey H: R^2^ = 0.69 and 0.79, respectively).

These dynamics in variance and autocorrelation are consistent with an underlying neuronal process that reflects accumulation, and are not easily explained by alternative accounts of LIP ramping activity such as a gradual shift of attention or simple movement preparation. Critically, the patterns were present over the same time window in both the high- and low-bet preferring populations. This appears to refute a version of the serial model where choice is initially resolved by considering only one pair of targets, followed by a shift to the other pair after some time has elapsed. We explicitly tested this by computing the expected autocorrelation under a simulated process where accumulation is delayed by a random amount of time. The delayed process provided a qualitatively inferior account of the empirically derived CorCE values, relative to standard (synchronous) accumulation (Fig. 4e, left vs. right; monkey H: p < 0.008, monkey G: p < 0.012 for both left-high and left-low populations). Taken together, the results support a parallel model in which deliberation occurs simultaneously between both the high and low pairs of targets. What remains to be tested is whether and when these accumulation signals are predictive of the monkey’s choice and wager on individual trials.

### Single-trial decoding reveals links between choice and confidence signals

Thus far most of our analyses have relied on trial averages, potentially obscuring the dynamics of individual decisions. We therefore turned to a population decoding approach (Kiani et al., 2014b; Kaufman et al., 2015; Peixoto et al., 2021; Steinemann et al., 2024; Charlton & Goris, 2024) to more directly address the question of parallel vs. serial deliberation. We trained two logistic classifiers, one for the binary choice and another for the binary wager, using the population spike counts (mean = 14 units/session) in the final 200 ms before the saccade. We then extracted a ‘neural decision variable’, also referred to as prediction strength or certainty (Kiani et al., 2014b; Peixoto et al., 2021), which is simply the log odds of a particular choice or wager as a function of time based on the decoded population spike counts on a given trial.

For both monkeys, the neural DV for choice ramped up starting about 200 ms after motion onset (Fig. 5a). The DV dynamics differed for the two animals, but both exhibited a ramping slope that depended on motion strength (monkey H: p < 0.001, monkey G: p < 10^−4^, linear regression). Cross-validated prediction accuracy also ramped up starting at this time, simultaneously for both the choice and confidence decoders (Fig. 5b). At their peaks, both performed well above chance on the test set, but a notable difference is the timing of the peaks, which for choice is just before saccade onset and for wager is slightly after the saccade (Fig. 5b). The time course of the choice and confidence DVs (Fig. 5c) mirrored the prediction accuracy traces, ramping in lockstep throughout most of the trial but with a subtle offset near the time of the saccade. This implies the persistence of a confidence-related signal even after commitment to a wager, possibly reflecting continued deliberation (or a top-down signal) which could drive CoMs or even inform the next decision (Lak et al., 2020). However, the offset was absent when using an alternative decoding approach with weights from a fixed (peri-saccadic) window (Supplementary Fig. 2), so this result should be interpreted with caution. The choice of method did not affect the main result of temporal congruency in the ramping of choice and wager signals during motion viewing.

**Figure 5.**
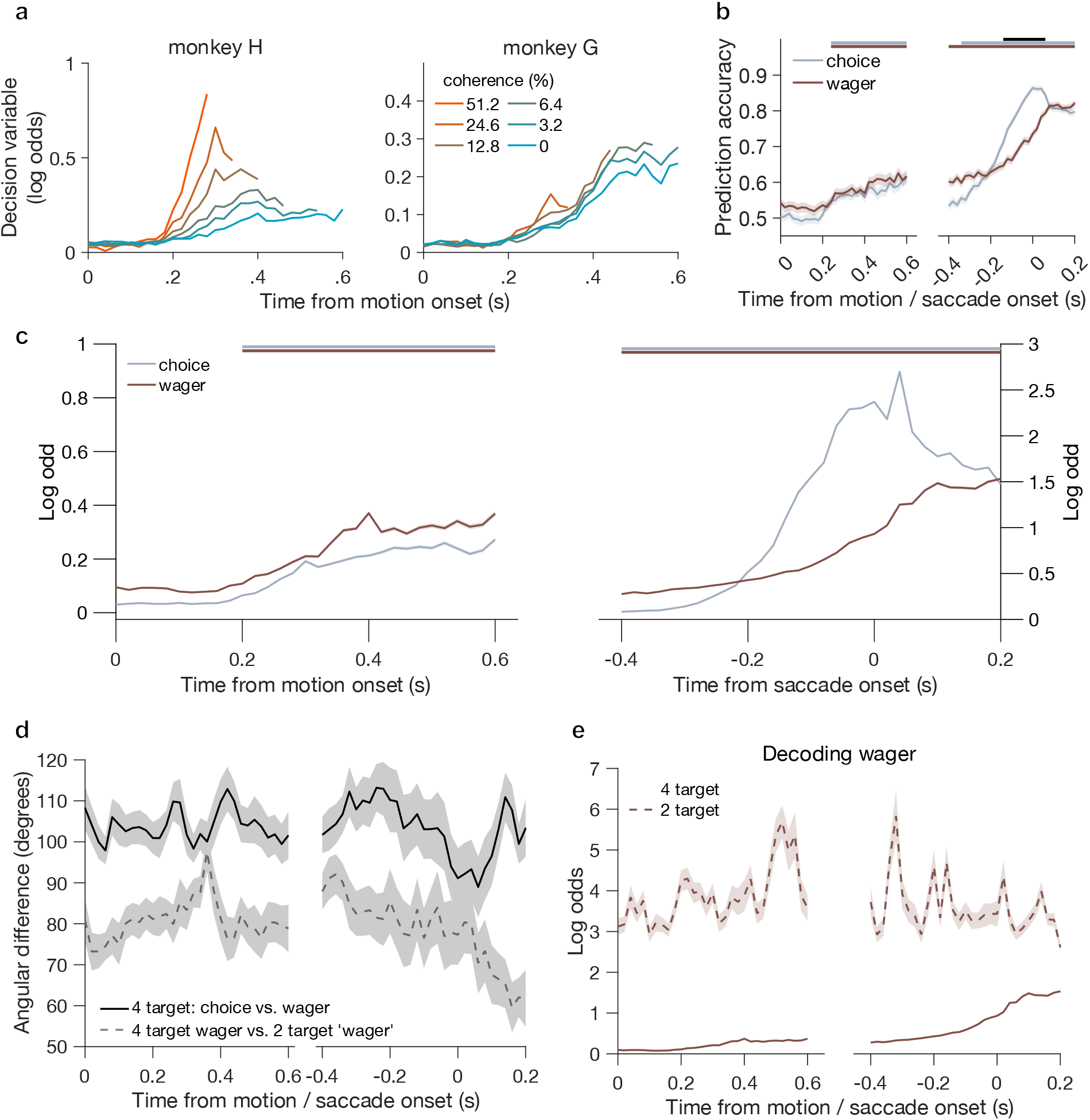
Population decoding of LIP activity supports concurrent, but orthogonal, readout of choice and wager. **a**, Log odds (‘neural decision variable’) quantifying prediction strength for the choice decoder as a function of time, aligned on motion onset and conditioned on motion strength. Left graph is monkey H and right graph is monkey G. **b**, Prediction accuracy (proportion correct binary classification in the test set), for both the choice (gray) and wager (brown) decoders, as a function of time and aligned to motion onset and saccade onset. Shaded regions around the traces indicate SEM. Data are from both monkeys. Gray and brown bars at top indicate when accuracy for the corresponding decoder was significantly greater than chance. Black bar indicates when prediction accuracy was significantly different for choice vs. wager. **c**, Log odds for choice and wager decoders as a function of time and aligned to motion onset and saccade onset. Line color, error shading, and significance bars are similar to **b. d**, Angular difference between the indicated pairs of decoder weight vectors as a function of time. Black solid = choice vs. wager on 4-target trials. Light grey = wager on 4-target vs. ‘wager’ on 2-target trials (vertical saccade component, collapsed across high-only and low-only control trials). Shaded regions are the SEM. For all three relationships the weight vectors appear closer to orthogonal than parallel. **e**, Wager decoder log odds as a function of time on trials with the standard 4-target configuration (solid trace, same as Extended Data Fig. 6b, wager) compared to 2-target control trials (dashed trace). Shaded regions indicate SEM.

Stepping back from questions of temporal alignment, we also examined the distribution of choice and confidence signals across the population, specifically asking whether these signals are represented by distinct subsets of neurons. We computed the correlation between the fitted decoder weights (choice vs. wager), after converting them to an absolute magnitude, and found a modest but highly significant correlation across the population (Extended Data Fig. 5a; monkey H: *r* = 0.18 & monkey G: *r* = 0.21; p << 0.05 for both, permutation test). Additionally, the distribution of the difference between choice and confidence weights was unimodal (Extended Data Fig. 5b; Hartigan’s dip test, p > 0.9 for both monkeys individually), suggesting a continuum of contributions to choice and confidence and not two distinct subpopulations. This raises the question of whether the population is capable of disentangling the two signals so that they do not interfere, especially given that the evidence that informs choice and confidence derives from a single source (the motion stimulus). To address this, we calculated the angular distance between the n-dimensional (n = number of neurons) decoding vectors for choice vs. confidence, separately for each session. During the deliberation period these vectors were approximately orthogonal (Fig. 5d, solid trace), facilitating the readout of confidence by a downstream region, potentially at any time—although they were closest to orthogonal around the time of the saccade. Lastly, to test whether concurrent choice and confidence signals could be a trivial consequence of motor preparation, we trained a separate decoder to predict the vertical component of the saccade using a set of control trials where only one pair of wager targets was present (either the high or the low). This decoding axis was nearly orthogonal to the vector for decoding the wager on standard 4-target trials (Fig. 5d, dashed trace), and showed a qualitatively different log odds profile (Fig. 5e), suggesting that the confidence signal is distinct from eye movement preparation in the absence of a wager decision.

As mentioned above, one can interpret the neural DV as a graded level of certainty (Kiani et al., 2014b) or even ‘confidence’ (Peixoto et al., 2019) in the prediction of choice by neural activity. Although such labels need not imply a neural representation of the corresponding probabilistic quantity (Pouget et al., 2016), we wondered whether the strength of choice decoding might predict the binary classification by the wager decoder on a trial-by-trial basis. To test this, we partitioned trials according to whether the wager decoder predicted a high or low bet (P(High) in the peri-saccade epoch greater or less than 0.5, respectively). We then averaged the DV from the choice decoder, using only 0% coherence trials, and found that it was higher for decoded-high vs. decoded-low trials (Fig. 6a). This indicates that the strength with which neural activity predicts the upcoming choice covaries with the probability that the same population predicts a high bet, consistent with a tight functional link between choice and confidence signals in LIP.

**Figure 6.**
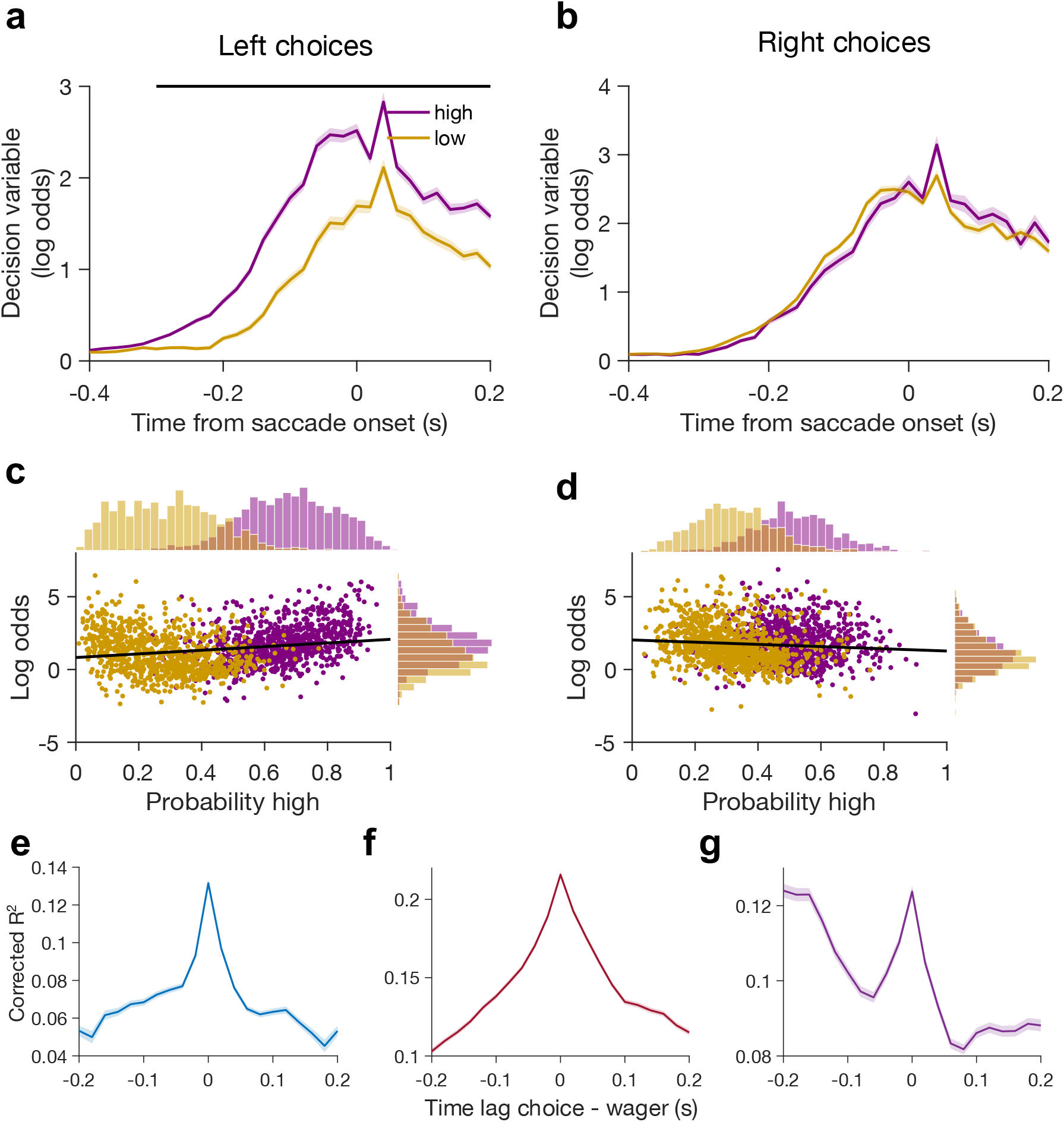
Predictive relationship between choice and confidence signals in LIP. **a**, Neural DV from the choice decoder as a function of time, aligned to saccade onset for leftward (contralateral) choices and 0% coherence trials only. Traces are separated by whether the wager decoder predicted a high (purple) or low (gold) bet. Shaded regions are SEM. Black bar at top indicates a one-way significant difference between the traces. **b**, Same as **a** but for rightward (ipsilateral) choice trials. **c**, Log odds from the choice decoder (leftward choices only) as a function of the probability of a high bet predicted by the wager decoder, based on the time window from −0.2s to 0.1s relative to the saccade. Each dot is an individual trial and the black line is a linear regression. Unlike A, purple and gold represents the behavioral wager outcome and not the decoder prediction. **d**, Same as **c** but for rightward (ipsilateral) choice trials. **e**, Corrected R^2^ values from a linear regression relating trial-by-trial choice decoding strength and wager decoder probability (P(High)), as a function of time lag between them. Values are computed using the time window 0 to 0.4 s from motion onset. Standard error indicated by the shaded regions (barely visible). **f**, Same as **e** but using the window −0.4 to 0 s relative to saccade onset. **g**, Same as **e** but using the window −0.2 to 0.2 s from saccade onset.

Remarkably, this link was only present for leftward (contralateral) and not rightward (ipsilateral) choices (Fig. 6a vs. 6b). To further investigate this stark contrast, we performed a trial-by-trial analysis of the neural DVs centered around the saccade epoch (Fig. 6c & 6d). After separating the data by the monkey’s wager on each trial (high=purple, low=gold), we confirmed that the wager decoder strongly predicted the behavioral confidence report, irrespective of choice (Fig. 6c & 6d top inset; p < 10^−250^, T = 56.5160, CI = [3.554 .3809] & p = 1.139×10^−188^, T = 33, CI = [0.1769 0.1993]). However, the wager prediction was only positively correlated with choice strength for leftward choices (Spearman rank correlation, *ρ* = 0.22, p =5.287×10^−22^ and *ρ* = −0.093, p=5.212×10^−5^ for left and rightward choices, respectively). Because we recorded only from the right hemisphere, the population is principally made up of neurons whose RFs overlapped a left choice target (either high or low). Thus, neurons which represent the unchosen option (associated with the ‘losing accumulator’ in the model) do not show a relationship between choice strength and wager prediction, even though they predict the wager itself (Fig. 6d, top histograms). This does not weaken our primary conclusion but highlights a gap between behavioral models and their neural implementation (see Discussion).

Having established a link between choice decoding strength and wager prediction (at least for contralateral choices), we can now examine the details of this relationship at a finer time scale and revisit the temporal offset shown in Figures 5b & 5c. We fit a linear regression relating choice decoder strength at time *t* to decoded wager probability at time *t* + *Δt*, where *Δt* ranges from +/-200 ms. We found that, during the deliberation phase (200-600 ms after motion onset and 400-0 ms before saccade onset), the best prediction between choice strength and wager probability was at a time lag of zero (Fig. 6e-f, R^2^ = 0.132 & 0.216 for blue and red lines, respectively). Interestingly, the period centered around the saccade (−0.2<->0.2s, purple line) gave rise to two peaks, one at zero lag (R^2^ = 0.124) and one at a lag of −0.2 s (choice preceding wager; R^2^ = 0.125). We speculate that this late peak may indicate a reevaluation of evidence informing the wager, akin to a replay of the last few samples of evidence leveraged for the choice as a substitute for external input. Regardless, the main takeaway is the prominent peak at zero lag, consistent with near-simultaneous updating of internal representations guiding a decision and confidence judgment—a surprising result given the prevalence of serial bottlenecks in various cognitive tasks and processes (Pashler, 1994; Welford, 1952; Sigman & Dehaene, 2005; Kang et al., 2021).

## DISCUSSION

The neurophysiological basis of metacognitive computations has become more accessible in recent decades through the development of behavioral assays of confidence in nonhuman animals (Hampton, 2001; Kiani et al., 2009; Middlebrooks & Sommer, 2012; Smith et al., 2012; Kepecs & Mainen, 2012). A longstanding goal is to connect the rich literature on process models for confidence with their implementation at the level of neural populations and circuits. One approach considers how decision accuracy, speed, and confidence can be jointly explained within the dynamic framework of bounded evidence accumulation (Pleskac & Busemeyer, 2010; Kiani et al., 2014; Fetsch et al., 2014b; Hellmann et al., 2023), an idea presaged by Vickers’ balance-of-evidence hypothesis (Vickers, 1979). Such a framework is motivated by the critical role of response time in psychophysical theory and experiment (Luce, 1986) and its strong empirical link to confidence going back at least a century (Henmon, 1911).

Embracing a dynamic model still leaves open questions about the temporal relationship between choice and confidence computations. Several authors have emphasized post-decisional processing (Baranski & Petrusic, 1998; Desender et al., 2021b), often formalized by serial models in which evidence is integrated for confidence only after termination of the primary decision (Pleskac & Busemeyer, 2010; Moran et al., 2015; Herregods et al., 2024). This idea follows naturally from the definition of confidence as the estimated probability correct conditioned on a choice (Pouget et al., 2016), and it is sensible to exploit any additional information acquired (or generated internally) after initial commitment. However, extending the accumulation process comes with both temporal and cognitive costs (Drugowitsch et al., 2012), and there are fundamental advantages to updating a provisional degree of confidence during decision formation (Song et al., 2017; Khalvati et al., 2021). Most decisions evolve in the context of ongoing or planned actions, executed with a degree of vigor proportional to expected utility (Shadmehr et al., 2019), which in turn depends on predicted accuracy (i.e., confidence). Recent work also suggests a role for confidence in strategic modulation of the ongoing decision process, including adjusting the termination criteria (Balsdon et al., 2020) and adapting to rapid changes in evidence strength (Drugowitsch et al., 2014; Balsdon & Philiastides, 2024). Lastly, considering the role confidence plays in sequential and hierarchical decisions (van den Berg et al., 2016b; Lisi et al., 2020; Zylberberg, 2021), computing confidence in parallel with each decision should make such sequences more efficient (Subramanian et al., 2019).

Although our study was not designed to directly probe such uses for ‘online’ confidence, it adds to the growing evidence that such a representation is generated and available during decision formation (Gherman & Philiastides, 2015; Dotan et al., 2018; Balsdon et al., 2020, 2021; Xue et al., 2023; Balsdon & Philiastides, 2024; Dou et al., 2024). It is notable that this signal is present within the same sensorimotor populations engaged in decision formation, raising the possibility of a local mechanism for confidence-gated changes in choice bias (Lak et al., 2020) or the tuning of sensory weights during perceptual learning (Law & Gold, 2008; Drugowitsch et al., 2019). Confidence computations have also been extensively linked to prefrontal, orbitofrontal, and cingulate cortices (reviewed by Fleming, 2024), raising the possibility that confidence is ‘fed back’ to sensorimotor cortices like LIP. This could, for example, explain the subtle temporal offset in our choice and wager decoders near decision termination (Fig. 5b-c; Fig. 6g), but seems harder to square with their tight alignment throughout most of the trial (Fig. 6e-f). It would be interesting to connect these phenomena to recent work suggesting that a separate ‘confidence accumulator’ exerts control over the primary decision (or motor) accumulator (Balsdon et al., 2020; Balsdon & Philiastides, 2024). Other accounts suggest that confidence is derived from an estimate of decision reliability or ‘meta-uncertainty’ (Shekhar & Rahnev, 2021; Mamassian & de Gardelle, 2022; Boundy-Singer et al., 2022), or governed by an accumulation of evidence bearing on stimulus detectability or discriminability (Maniscalco et al., 2021; Hellmann et al., 2023). A combination of large-scale recordings and causal manipulations will likely be necessary to develop a mechanistic account that unifies these observations and explains why the brain computes such a diverse and anatomically distributed array of metacognitive signals.

We found that the strength of the decoded choice prediction was correlated with the probability of a high bet predicted by the wager decoder (Fig. 6a, 6c). This is important because it suggests a direct link between the DV representation and readout of confidence, as predicted by models instantiating a so-called ‘common mechanism’ (Kiani & Shadlen, 2009; Fetsch et al., 2014b; van den Berg et al., 2016a). However, the relationship held only for contralateral choices (Fig. 6b, 6d), i.e. for trials where the recorded neurons are presumed to represent the ‘winning’ accumulator in a race model. Superficially, this seems to contradict a basic tenet of such models (Kiani et al., 2014a), namely that the *losing* accumulator implements the mapping between accumulated evidence and probability correct, since the winner is always at the bound at the time of the decision. We considered an alternative behavioral model that reads out confidence from the winner (Le Denmat et al., 2024), but this failed to capture the differences in accuracy and RT conditioned on the wager (Supplementary Fig. 2), unless there is at least 50-60 ms of post-decision accumulation for confidence. However, this in turn predicted a prominent folded-X pattern (Supplementary Fig. 3f & 3l) that was absent in the data. How can we reconcile this apparent conflict between the modeling, favoring a losing-accumulator mechanism, and the neural evidence supporting a stronger link with the winner (see also ref. Zylberberg & Shadlen, 2025)? Note that our 4-target task presumably entails competition between four spatially defined neuronal pools, in contrast to the two competing accumulators of the model. This exemplifies the gap that generally exists between the level of explanation provided by psychological models and the implementation of a given confidence-guided behavior. Bridging this gap will require additional modeling and simulation, ideally constrained by the dynamics and correlation structure across subpopulations (So & Shadlen, 2022) and the heterogeneity of functional cell types that may be hidden in population averages (Zylberberg & Shadlen, 2025).

One limitation of the current work is that confidence is mapped onto a stable motor action, namely the saccade to a high or low target whose positions did not change within a session. We found this to be necessary for achieving consistent behavioral performance, but it does present interpretational challenges related to the overlap of cognitive and motor-planning signals. Could the results be explained solely by motor preparation to one of four independent spatial targets? We do not think so. First, the observed signatures of noisy evidence accumulation (Fig. 4e, Fig. 5a) suggest that ramping activity in LIP reflects more than mere saccade planning. Second, the population activity pattern predicting the wager was distinct from the pattern preceding the same saccade on control trials when only one pair of targets was available (Fig. 5d-e). This does not mean the results are incompatible with embodied or ‘intentional’ theories of decision making (Cisek, 2007; Shadlen et al., 2008); quite the opposite. We interpret the overlap of metacognitive and premotor signals as supporting and extending the idea of an intentional framework for decisions among actions, characterized by ‘continuous flow’ of information to the motor system (Selen et al., 2012). In this context, it is intriguing that two physical dimensions of a motor plan (horizontal and vertical) can be updated simultaneously based on distinct transformations of the input (a categorical judgment and prediction of accuracy). These were by no means guaranteed to be computed in parallel (Zylberberg et al., 2012b); in fact, a recent study of 2D decisions using a similar target configuration (simultaneous report of color and motion; Kang et al., 2021) suggests there is a bottleneck preventing simultaneous incorporation of two evidence streams into a single DV. Evidently there is no such bottleneck for a single evidence stream informing choice and confidence, at least as far as we can resolve with neural decoding.

Our study exposes one type of joint representation of choice and confidence in a sensorimotor region, yet even early visual cortical neurons carry information about uncertainty (Walker et al., 2020; Hénaff et al., 2020) and are predictive of confidence (Geurts et al., 2022; Boundy-Singer et al., 2024). A key question is how sensory activity is read out in a feedforward manner to update premotor and higher-order metacognitive representations. Conversely, feedback of ‘belief’ signals is a cornerstone of theories of perceptual inference (Lee & Mumford, 2003; Haefner et al., 2016), but whether this process is related to the explicit sense of decision confidence and its neural correlates is unclear. It seems likely that unifying the feedforward-feedback dichotomy will be necessary for uncovering general principles by which population dynamics establish internal belief states, while simultaneously generating adaptive behavior in uncertain and changing environments.

## Acknowledgements

We thank members of the lab for their insight and discussions. Additionally, we are grateful to Ofelia Garalde for animal assistance, and Justin Killebrew, Bill Nash, and Bill Quinlan for technical assistance. This work was supported by the National Institute of Neurological Disorders and Stroke (RF1NS132910), the E. Matilda Ziegler Foundation for the Blind, and a Whitehall Foundation research grant. CF is supported by the France-Merrick Foundation.

## Data Availability Statement

Data will be made available without restriction via the DANDI archive upon acceptance for publication.

## Code Availability Statement

All code needed to reproduce the figures and analyses will be made available upon request after acceptance for publication.

## METHODS

### SUBJECTS AND EXPERIMENTAL PROCEDURES

Two male rhesus monkeys (*Macaca mulatta*, 6-8 years old, 8-10 kilograms) were handled according to National Institutes of Health guidelines and the Institutional Animal Care and Use Committee at Johns Hopkins University. Standard sterile surgical procedures were performed to place a PEEK recording chamber (Rogue Research) and titanium head post under isoflurane anesthesia in a dedicated operating suite. The recording chamber was positioned over a craniotomy above the right posterior parietal cortex of both animals for access to the intraparietal sulcus and posterior third of the superior temporal sulcus. The chamber and head post were held in place using dental acrylic anchored with ceramic bone screws.

### Experimental apparatus

Monkeys were seated in a custom-built primate chair in a sound-insulated booth facing a visual display (ViewPixx, VPixx Technologies; resolution 1080×960, refresh rate 120 Hz; viewing distance 52 cm) and infrared video eye tracker (Eyelink 1000 Plus, SR Research). Experiments were controlled by a Linux PC running a modified version of the PLDAPS system (Version 4.1, Eastman & Huk, 2012) in MATLAB (The MathWorks). Visual stimuli were generated using Psychophysics Toolbox 3.0 (Brainard, 1997).

For correct responses, the monkey was given a fluid reward that was dispensed using a solenoid-gated system.

### Neurophysiology

Recording probes (32- or 128-channel Deep Array, Diagnostic Biochips) were positioned with the aid of a PEEK grid secured inside the recording chamber. A sharpened guide tube was inserted through a grid hole so that the tip of the tube just punctured the dura, then a probe was advanced through the guide tube into the brain using a motorized microdrive (40mm MEM Drive, Thomas Recording). Bandpass-filtered voltage signals were collected using the Open Ephys acquisition board and software (Siegle et al., 2017). Post-hoc analysis for identifying single neurons and multi-unit clusters was done using Kilosort 2.0 (Pachitariu et al., 2016; Stringer et al., 2019) with additional curation using phy2 software (https://phy.readthedocs.io/en/latest/). Data analysis was performed with custom MATLAB code.

Targeting of LIPv was achieved by selection of grid locations based on a post-surgical structural MRI scan in which the chamber and grid holes were well visualized. We compared the MR images to published reports and atlases (Lewis and Van Essen, 2000; Saleem & Logothetis, 2006) to estimate the depth of LIPv (typically 8-12 mm from the dura in our vertical penetration angle) and corroborated the targeting using white-gray matter transitions and physiological response properties during the mapping tasks described below. After reaching the target, we let the probe settle for 30-60 minutes before the start of the experiment. A total of 407 units (single = 148 & multi = 56 in monkey H; single = 107 & multi = 93 in monkey G) were collected over 29 sessions (12 for monkey H, 17 for monkey G). No qualitative differences were detected in the results using single vs. multi-units so they were pooled for all analyses.

### Memory saccade task

Sessions began with a standard memory-guided saccade task to identify neurons with spatially selective activity during the delay period (Gnadt & Andersen, 1998) and to coarsely map their receptive fields (RFs). Monkeys were instructed to gaze at a central fixation point (1.5° radius acceptance window), after which a red target (0.42° diameter circle) was flashed for 100 ms, located in one of several locations evenly spaced in polar coordinates. The coordinates consisted of 3 different radii (eccentricities) and 10 or 12 angular positions, giving a total of 30 or 36 unique target locations. Each target location was presented 10 times in pseudorandom order, requiring a total of 300 or 360 trials. While fixating, the monkey had to remember the location of the target, and after a delay of 0.8 s the fixation point was extinguished, instructing the monkey to make a saccade to the remembered location. RFs were estimated online during/after the memory saccade block by acquiring multi-unit spikes (threshold crossings) on each recording channel and plotting the mean firing rate during the memory delay as a function of target location in a 2D heat map. These RF maps guided the placement of the four targets for the main decision task, such that at least one of the targets overlapped the RF of multiple neurons in the recorded population. Neurons whose RFs did not overlap any target were excluded from further analysis.

### Main task

The monkeys were trained to perform a reaction-time direction discrimination task with simultaneous report of choice and confidence (‘peri-decision wagering’; Fig. 1a). To initiate a trial the animals acquired fixation on a target on the center of the screen (0.21° diameter). After a delay of 0.5 s four targets appeared, positioned diagonally from the center of the screen, each representing a choice (left or right) and a wager, or bet (high or low). The targets representing *high* (low) bets were always placed in the upper (lower) quadrants (mean ± S.D. of [x,y] target positions relative to fixation: left-high = [−7.3±2.1°, 7.3±1.5°]; right-*high* = [7.3±2.1°, 7.3±1.5°]; left-low = [−6.9±2.3°, −3.4±1.4°]; right-low = [6.9±2.3°, −3.4±1.4°]. Each left-right pair was presented symmetrically around the vertical meridian, but high-bet targets were typically 2-5° more eccentric than low-bet targets in order to counteract the monkeys’ tendency to bet high more often than low.

After another brief delay (0.3-0.6 s, truncated exponential), a dynamic random-dot motion (RDM) stimulus was presented in a circular aperture. Motion strength, or coherence, was sampled uniformly on each trial from the set {0%, ±3.2%, ±6.4%, ±12.8%, ±25.6%, ±51.2%}, where positive is rightward and negative is leftward. The stimulus was constructed as three independent sets of dots (Roitman & Shadlen, 2002), each appearing for a given video frame then reappearing three frames (25 ms) later. Upon reappearing, a given dot was either repositioned horizontally to generate apparent motion in the assigned direction (speed = 2-16°/s, held constant within a session) with probability given by the coherence on that trial, or otherwise was replotted randomly within the aperture.

When ready with a decision, the animal could report its choice and wager by making a single saccade to one of the four targets. When the eyes moved 1.5° away from the target the RDM and fixation point were extinguished while the four targets remained visible. When the eye position reached one of the four targets, it was required to hold fixation within 1.5° of the target for 0.1 s to confirm the outcome. Lastly, the animal was either rewarded or given a time penalty depending on the conjunction of accuracy (choice corresponding to the sign of coherence) and wager (Fig. 1a, right). The penalty for a high-bet error was applied to the subsequent trial where the animal was required to fixate the central target for a longer period of time (2-3 s) prior to RDM onset. Reward sizes (~0.21 ml for high and ~0.19 ml for low bets) and penalty times were chosen to encourage a wide range of wager frequency across different levels of motion strength.

### QUANTIFICATION AND STATISTICAL ANALYSIS

#### Cell selection

For consistency with previous work, we limited the analyses shown in Fig. 4 and Extended Data Fig. 4 to units with spatially-selective persistent activity, based on the memory saccade task. We quantified spatial selectivity using a previously described discrimination index (Nguyenkim & DeAngelis, 2003):

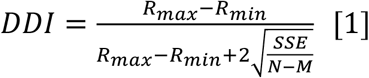

where *R*_*max*_ and *R*_*min*_ are the mean firing rates during the delay period at the target location with the highest and lowest response, respectively. *SSE* is the sum-squared error around the mean responses, N is the total number of trials, and M is the total number of unique target locations. For Fig. 4 and Extended Data Figs. X (all analyses except the population logistic decoder), only units with DDI > 0.45 were included (N=195). This criterion is somewhat arbitrary but was based on qualitative inspection of numerous individual RF maps, and chosen prior to designing and executing the main analyses.

For assigning units a preferred target location in the peri-dw task, we quantified RFs offline by fitting a 2D Gaussian to the firing rates during the delay period of the memory saccade task:

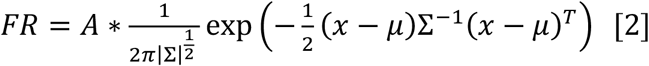

where FR is firing rate during the delay period and x is a 2-length vector that contains the X and Y coordinates of the memory saccade targets. The fitted parameters include *A* for amplitude, *μ* for the position of the RF center (X and Y in degrees), and *Σ* which is the 2×2 covariance matrix of the Gaussian. To reduce the variables for fitting we set the covariance to 0 and only fit for the two variances. We then normalized the 2D Gaussian to convert it to a probability density and defined a unit’s preferred location based on which of the 4 choice-wager targets was associated with the highest probability.

For the population logistic decoders, we included all well-defined single neurons or multi-unit clusters based on careful spike sorting and manual curation. Previous studies have suggested that decision-related activity can be present even in neurons without spatially-selective persistent activity (Meister et al., 2013), and we aimed to maximize the sample size to facilitate single-trial decoding. This broader criterion increased the average number of units per session from 7.5 to 17.3 for monkey H and from 6.2 to 11.2 for monkey G.

#### Behavioral data analysis

We applied a logistic regression model to fit the proportion of rightward choices:

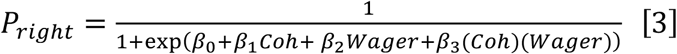

where *P*_*right*_ is the probability of a rightward choice, *Coh* is signed motion coherence, *Wager* is the monkey’s bet (high/low), *β*_0_ is the overall bias, *β*_1_ estimates choice sensitivity, *β*_2_ captures any bias related to the bet (typically zero), and *β*_3_ is the interaction term used to test whether choice sensitivity depends on wager. Fitting was done by minimizing the negative log-likelihood under a binomial distribution, using *fminsearch* in MATLAB with the Nelder-Mead method.

We used a modified Gaussian function to provide descriptive fits of the mean reaction time (RT) as a function of motion strength, as follows:

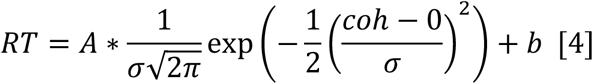

where *A* is an amplitude term, *coh* is the signed motion coherence, *σ* is the standard deviation controlling the width of the Gaussian, and *b* is a baseline term capturing the fastest mean RT. We fit this pseudo-Gaussian by minimizing root-mean-square error (RMSE).

To examine the relationship between accuracy and RT, as well as wager and RT, we calculated proportion correct and proportion of high bets grouped by RT, using non-overlapping 100 ms time bins starting 100 ms after motion onset. To test for significance as to whether the trend was decreasing (during the time depicted in Fig. 1d) we used a Cochran-Armitage test for trend:

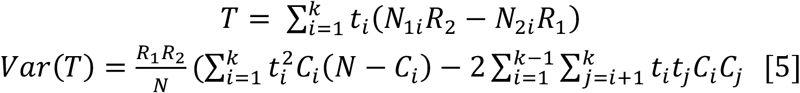

where *t*_*i*_ are the weights depicting the trend (in our case linear, so t = [0, 1, 2, 3, 4, …]), *N* = total number of trials, *N*_*ab*_ is total number of trials for group *a* (accuracy: correct and error; wager: high or low) and *b* is total number of trials for timepoint *b. R*_*a*_ represents total number of trials for group *a* irrespective of time, and *C*_*b*_ is total number of trials at timepoint *b* irrespective of group. The division of *T* by *Var(T)* gives a test statistic that can then be used to compute a p-value.

#### Motion energy analysis

To estimate the temporal weighting of sensory evidence for choice and confidence, we utilized motion stimulus fluctuations to perform a psychophysical reverse correlation analysis on both choice and wager. We convolved each trial’s sequence of dots — a 3-dimensional array with the first two dimensions denoting the X-Y coordinates of the dots’ center, and the third dimension spanning the number of frames — with two pairs of quadrature spatiotemporal filters (Adelson & Bergen, 1985). The filters were oriented to account for motion in the direction along the choice axis: 0° (rightward) and 180° (leftward). The convolved quadrature pairs were squared and summed to give the local motion energy for both the leftward and rightward directions. These local motion energies were then collapsed across space (first 2 dimensions) to derive the rightward and leftward motion energy provided by the stimulus through time, which we then subtracted (right minus left) to obtain the net motion energy (Adelson & Bergen, 1985; Kiani et al., 2008). To strictly look at the fluctuations around the mean and mitigate potential effects of coherence, we subtracted from each trial the mean motion energy, conditioned on signed coherence, through time. As most meaningful fluctuations occur at low coherences, we also restricted the analysis to 0%, 3.2% and 6.4% coherence trials.

For the choice kernels (Fig. 2a-b) we simply averaged the net motion energy time series for all trials with a right or left choice. To evaluate statistical significance we used a t-test comparing the motion energy profiles for right and left choices at each time point, applying a Šidák correction matching the number of time samples. For the confidence kernels (Fig. 2c-d), we instead subtracted the motion energy profiles for high vs. low wager trials conditioned on a given choice (correct trials only) and computed the standard error of the difference (shaded area around traces in bottom of Fig. 2c-d). To assess significance, we compared the confidence kernel distribution relative to a value of zero using a t-test with Šidák correction for the number of time samples.

#### Changes of mind (CoMs) and changes of confidence

Raw eye position data (sampled at 1000 Hz) were converted to velocity and smoothed by applying a 3rd-order low-pass Butterworth filter with a cutoff frequency of 75Hz (Orozco et al., 2021). Eye position is a 2D vector containing X and Y position in degrees, while eye velocity was defined as 1D vector that combines the velocities for both directions (X and Y). To preprocess the data, we first calculated a stricter time for saccade onset by applying a threshold of 20°/s onto the smoothed velocity data. Subsequently, we centered the eye positions by subtracting the average of the last 5 ms before saccade onset. The initial choice and wager were defined by which quadrant of the screen the eyes were located in 5 ms after saccade detection. The final choice and wager corresponded to the target at which the eye position settled within a 0.1 s grace period following the initial saccade. This provided an initial and final choice and wager for every trial, allowing for simple analyses like those for Figure 2e-j. CoM frequencies (Fig. 2e-f, 2h-i) were conditioned on the initial outcome, therefore the frequencies reflect not the proportion of all trials but only trials that initially reached a given outcome (error, correct, low, and high). For Figures 2g & 2j the probabilities are reflective of all trials, hence the values are much smaller than on the other plots.

#### Parallel model

To formalize the hypothesis of parallel deliberation for choice and confidence, we used a two-dimensional (2D) bounded accumulator model (Kiani et al. 2014), also known as an anticorrelated race. We adapted a recently developed family of closed-form solutions for a 2D correlated diffusion process (Shan et al., 2019) to facilitate fitting of the parameters. By conceptualizing the diffusion process as a Gaussian distribution originating from the third quadrant on a plane with two absorbing bounds, one can employ the method of images (MoI) to calculate the propagation of the probability density of the diffusing particle, i.e. the solution to the Fokker-Planck equation. The constraint making this numerical solution possible limits the discrete number of anticorrelation values that can be modeled, governed by the number of images:

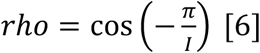

where *rho* is the correlation value and *I* is the number of images. We selected *I = 4* or *rho* = .7071 for consistency with previous studies (Kiani et al., 2014a; Van Den Berg et al., 2016a).

Specifically, the MoI yields *P*(*v*_*right*,_ *v*_*left*_/*C*,*t*), describing the probability of the accumulator being in a particular position at time *t* for coherence *C*. The probability of making a right choice is given by:

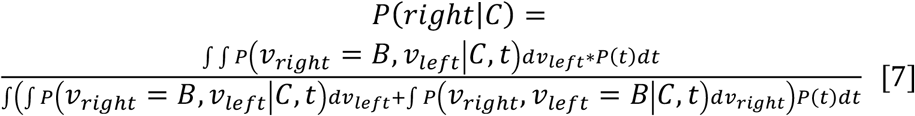

where *B* is the bound for terminating the accumulation process. To obtain the decision time (DT) distribution, we calculated the difference in the survival probability as follows:

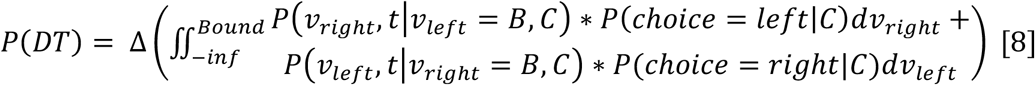

thereby providing the change in probability at each survival timestep, i.e. the probability of crossing a bound. The reaction time (RT) is the decision time plus sensory and motor delays, referred to as non-decision time or *nonDT*, and is obtained by convolution as follows:

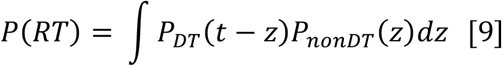

where *P*_*nonDT*_*(z)* is modeled as a Gaussian distribution with mean *(μ*_*nonDT*_) and standard deviation *(P*_*nonDT*_).

To calculate the probability of betting high, we first computed the log odds of a correct choice as a function of the state of the losing accumulator (Kiani et al., 2014), as follows:

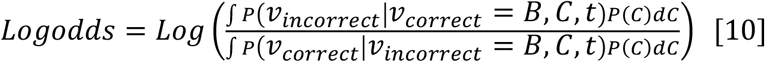

where *v*_*incorrect*_ is the ‘incorrect’ accumulator (not matching the sign of coherence) and *v*_*correct*_ is the correct accumulator (matching the sign of coherence). This transformation provides a graded scale for confidence that can be transformed into {high, low} wager responses by applying a cutoff value (*θ*) that imposes binary outcomes. To obtain the probability of a high bet, we computed:

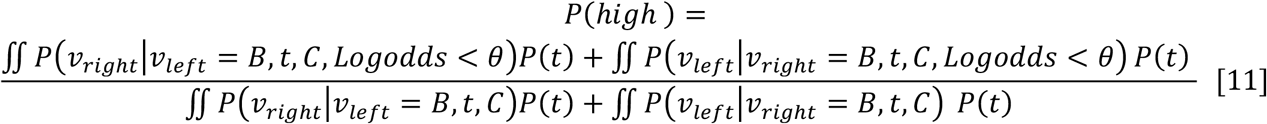

where *Logodds < θ* indicates integration over the area in which *Logodds* is less than the cutoff value.

We found that the fits were improved with two subtle changes from the model used previously (Kiani et al., 2014a; van den Berg et al., 2016a), although they did not affect the conclusions regarding serial vs. parallel. The first is an extra parameter (τ_*cut*_) that acted as a cutoff marking when confidence no longer depended on elapsed time but only the amount of evidence accumulated. This relaxes the assumption of an optimal mapping from DV to confidence, which is time-dependent due to marginalization over a mixture of motion strengths (Kiani & Shadlen, 2009). Second, having observed occasional changes of mind (Fig. 2e-j), we considered the possibility that the brief post-decision epoch that mediates CoMs also permits some temporal flexibility in the assignment of confidence. Specifically, we found that the wager-conditioned RT distributions were better fit when conditioning on the wager that would have been made at the end of the non-decision time, rather than at decision time. This result defies simple interpretation under the current modeling framework, but it may point toward an interesting target for more complex models in future work.

We obtained the best-fitting parameters to the model by using the joint probability of choice, RT, and wager as follows:

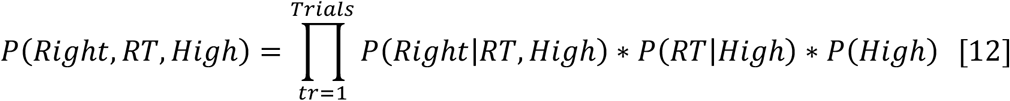

We calculated the negative log likelihood of this by:

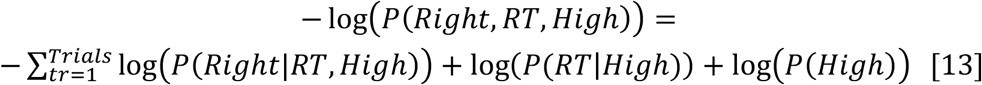

The likelihood calculation thus optimizes for choice and RT conditioned on wager (Fig. 3, left and middle columnd) but the wager itself is unconditioned, such that the split between correct and error trials in Fig. 3 (right column, green vs. purple) is a prediction, not a fit. The full model had 8 or 7 free parameters, differing slightly between the two animals (see table below). These included drift rate *(K)*, bound *(B)*, mean non-decision time (*μ*_*nonDT*_), confidence time cutoff (τ_*cut*_), wager offset (*b*_*w*_), and log odds cutoff *(θ)*. Monkey H required separate non-decision time means for left and right choices, and a ‘wager offset’ (*b*_*w*_) capturing a tendency to occasionally bet low across the board, even for the highest coherences (akin to a lapse rate). In contrast, monkey G tended to ‘lapse’ (bet low more often than expected) only for weaker motion strengths, requiring an ad-hoc gain factor applied to the *P*(*High*) distribution in a coherence-dependent manner (parameterized by *gain* = 1 + α * (1 − ϵ * | *coherence*|).

Non-decision time standard deviation (*σ*_*nonDT*_) was not a free parameter and was instead established using psychophysical kernels (van den Berg et al., 2016a). Fitting was done using MATLAB’s built-in function *fminsearch* applied with a grid-search method with 30 different starting points.

| Monkey | Parameters | Number of Parameters |
| --- | --- | --- |
| Hanzo | $K, B, \mu_{nonDT, coh+}, \mu_{nonDT, coh-}, \theta, b_{w, coh+}, b_{w, coh-}, \tau_{cut}$ | 8 |
| Genji | $K, B, \mu_{nonDT}, \theta, \tau_{cut}, \alpha, \epsilon$ | 7 |

#### Serial model

The serial model was constructed as a sequence of two 1D bounded accumulator (drift-diffusion) models, one for choice followed by a second for confidence (Fig. 1f). There are 6 main parameters: drift rate *(K)*, choice bound *(B*_*c*_), high wager bound *(B*_*h*_), low wager bound (*B*_*1*_), mean non-decision time (*μ*_*nonDT*_), and a linear urgency signal for only the confidence accumulator (*u*_*M,Conf*_). Parameter estimation largely followed the same logic as the parallel model, using the joint distribution of choice, RT, and wager to fit the data, with a few minor differences. Computing the probability of rightward choice was similar to the parallel model and used the following formula:

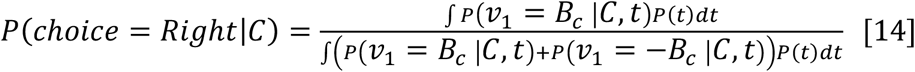

where *ν* is the decision variable for the first 1D accumulator, and the other parameters are similar to the parallel model. The decision time distribution was calculated as:

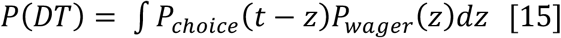

where *P*_*choice*_ is the probability of hitting a bound at time t for the first accumulator and *P*_*wager*_ is the probability of hitting a bound at time t for the second accumulator. For RT we followed the same procedure as the previous model. Lastly, to calculate the probability of betting high, we used the following equation:

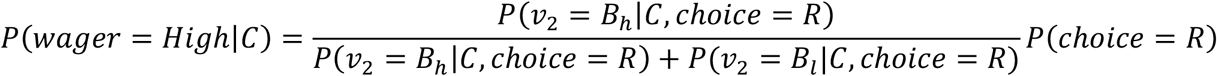

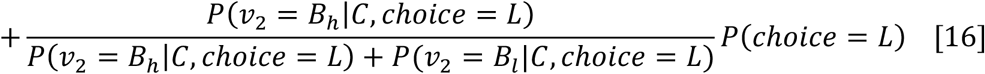

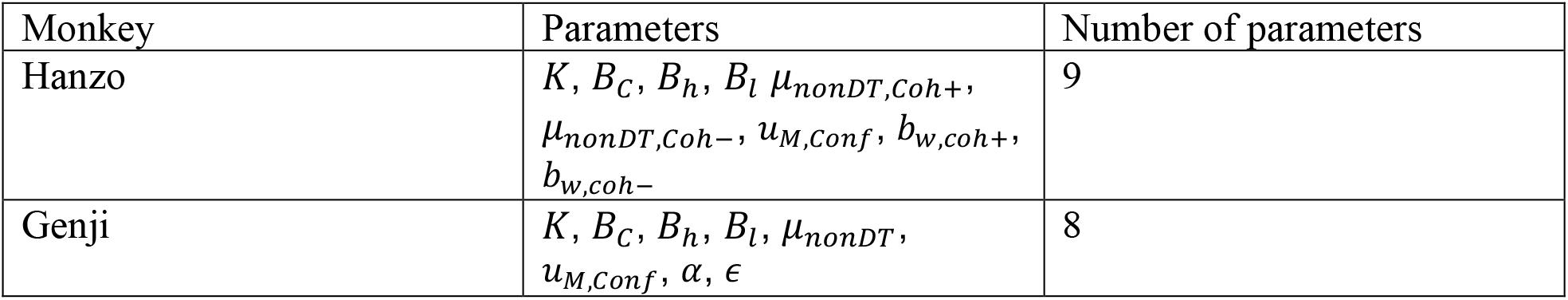

#### Hybrid model

The hybrid model was formalized as a two-dimensional race, thereby also using the closed-form solution (Shan et al., 2019). Given the similarity to the parallel model, all variables were computed using the same formulae, except for an additional convolution step to compute the distribution of decision time plus the extra time used for the wager. We defined ‘wager time’ (WT) as the period that the losing accumulator is allowed to continue after the winning accumulator reaches its bound. The total decision time is formulated as:

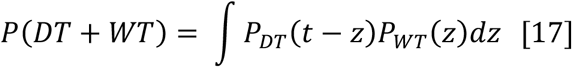

where DT represents the time for both choice and wager and WT strictly the extra time for confidence. Reaction time is then computed by convolving *P(DT + WT)* with the non-decision time, as in Equation 9. Subsequent computation of P(wager) and its dependencies are computed using *P(DT + WT)* instead of *P(DT)*.

| Monkey | Parameters | Number of Parameters |
| --- | --- | --- |
| Hanzo | $K, B, \mu_{nonDT, coh+}, \mu_{nonDT, coh-}, \theta, b_{w, coh+}, b_{w, coh-}, \tau_{cut, coh+}, WT$ | 9 |
| Genji | $K, B, \mu_{nonDT}, \theta, \tau_{cut}, \alpha, \epsilon, WT$ | 8 |

#### Analysis of neural data based on preferred and chosen target

Two populations of neurons were established based on RF overlap of either the left-high or left-low target. We computed four normalized firing rate responses for both populations of neurons, where each response corresponds to one of the four chosen targets (Fig. 4b-c). To combine across neurons, we first detrended the responses of each neuron over the coherences of interest [−6.4%, −3.2%, 0%, 3.2%, 6.4%]. Then the firing rates for each neuron (after smoothing each trial with a 0.1 s exponential filter) were averaged over all trials that met 3 conditions: 1) matched the chosen target of interest, 2) contained a coherence of interest, and 3) had a RT longer than 0.3 s. Example units in Figure 4a were not detrended or normalized but otherwise the procedure was the same. The colored bars at the top of Figures 4b-c indicate statistical significance based on a one-tailed t-test evaluating whether the activity preceding a choice of the RF-aligned target was greater than each of the other three targets, indicated by the color. Given that the testing is done over multiple timepoints, the one-tailed t-test alpha value was adjusted using Šidák’s correction.

#### Signatures of accumulator dynamics in firing rate variance and autocorrelation

An accumulation of noisy evidence produces characteristic variance and autocorrelation features that can be estimated from single neurons using procedures laid out by previous work (Churchland et al., 2011; de Lafuente et al., 2015; Shushruth et al., 2018; Steinemann et al., 2024). Applying the law of total variance to a doubly stochastic process, the variance in spike rate for a given time bin is a summation of the variance of the underlying latent rate, termed variance of the conditional expectation (VarCE), and the residual variance expected if the latent rate was constant, known as point process variance (PPV). To calculate VarCE, one must subtract out the PPV from the total measured variance. To do this we made two simple assumptions (Churchland et al., 2011): (1) the observed spiking of a neuron follows a stochastic point process mediated by some rate parameter, and (2) at each time bin the PPV is proportional to the mean count:

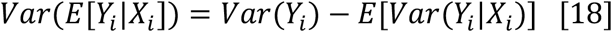

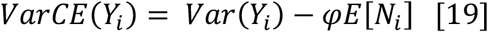

where *Y*_*i*_ represents the random variable capturing the neuron’s spike count at timepoint *i*,*X*_*i*_ is the random variable for the latent rate at timepoint *i*, and *φ* is a constant that is fitted to maximize how well the observed firing rates match an accumulation of independent identically distributed (iid) random numbers. *E*[*Y*_*i*_] is the mean spike count at timepoint *i*. In addition, it follows that the law of total covariance is described using a similar equation:

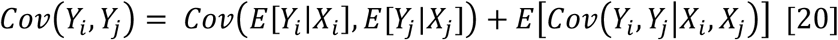

The first term on the right-hand side is known as the covariance of conditional expectations (CovCE), which is needed to compute the correlation of conditional expectations (CorCE). The second term is the expectation of conditional covariance, and its diagonal is the PPV. To calculate the CorCE, we made another assumption that when *i* ≠ *j*, the expectation of conditional covariance is zero because the variance from the point process should be independent across timepoints (Churchland et al., 2011; although this may not be strictly true for adjacent time bins due to their shared interspike interval). This simplification makes it possible to state that the CovCE, for *i* ≠ *j*, is equal to the measured covariance, and the diagonal of the CovCE is then the VarCE. It follows that to calculate the CorCE, one must simply divide the CovCE by the VarCE:

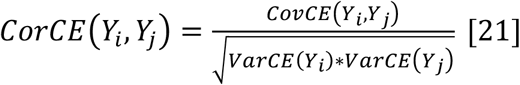

where *i* and *j* are time points. The best-fitting *φ* is then calculated by comparing the empirical CorCE estimates with theoretical or simulated correlation values under the hypothesized generative process (e.g., accumulation of iid samples).

We tested two theoretical autocorrelation patterns, one pertaining to a standard drift-diffusion process and the other to a delayed drift-diffusion process (Fig. 4d). The standard accumulation of iid random numbers was calculated using the following equation:

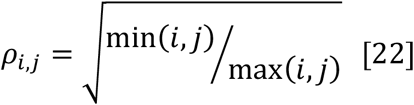

We used 6 different timepoints giving 15 unique combinations *(i = 1*: 6 & *j = j*: 6, *i* ≠ *j)*. For the delayed accumulation of iid random numbers, we used a simulation that accumulated noisy normalized samples of numbers with mean [0.717, 0, −0.717]. We narrowed the simulation to the first 6 timesteps to compare it to the results from the standard accumulation process. Additionally, the delay component was constructed by uniformly sampling a value between 1-6, indicating when the accumulation would begin. Using 10,000 trials we calculated the autocorrelation of the first 6 timesteps for this simulation providing 15 unique combinations *(i = j*: 6 & *j = j*: 6, *i* ≠ *j)*. Additionally, we fit the *φ*, for both models, according to the following steps: (1) calculate the *E*[*Y*_*i*_], *Var(Y*_*i*_), and *Cov*(*Y*_*i*_,*Y*_*j*_) from observed spikes; (2) compute *Var CE*(*Y*_*i*_) using an initial value of *φ* = 1; (3) calculate *CorCE*(*Y*_*i*_,*Y*_*j*_) under the assumptions mentioned above; (4) calculate the mean squared error (MSE) between the empirical *CorCE(Y*_*i*_, *Y*_*j*_) and the theoretical/simulated autocorrelation values, *ρ*_*i*,*j*_; and (5) iteratively update *φ* until the MSE between the *CorCE(Y*_*i*_, *Y*_*j*_) and *ρ*_*i*,*j*_ reached the global minimum.

We used 6×60-ms time bins spanning from 170 ms after motion onset to 530 ms after motion onset. We applied the analysis on trials with coherences of [−6.4%, −3.2%, 0%, 3.2%, 6.4%] and reaction times at least 630 ms to minimize bound effects. To combine across neurons, we calculated the mean response for each time bin of each neuron across all trials and subtracted that from the mean response for each time bin conditioned on the signed coherence. This then gives a matrix of residuals that is of size [*neuron * trial x time bins*], which is then used to calculate a covariance matrix. Next, the VarCE is calculated by substituting the raw variance for the diagonal in the covariance matrix, since the diagonal is the normalized population variance, and *φ* is initiated at a value of 1. To calculate the empirical correlation values, each entry in the covariance matrix is divided by 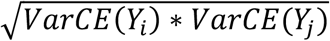. Lastly, using a fitting procedure we compare the Fisher’s z-transformation of the empirical correlation with a z-transformation of the ideal correlation and minimize the mean squared error (MSE) of the correlation matrix.

Testing for significant differences between standard and delayed accumulation was done using a leave-one-out (LOO) cross-validation method. The metric used to test the validation was the mean absolute percentage error (MAPE). MAPE allowed for comparison between the two models because both models contained different dependent variables (*ρ*_*i*,*j*_). This method provided 15 different MAPE values for each model, which were then compared using a one-tailed t-test. The model with the lowest percentage error distribution better captured the underlying autocorrelation structure of the data (Fig. 4e).

For Extended Fig. 4 we recalculated the VarCE but used 6×100-ms time bins spanning −50 ms before motion onset to 550 ms after motion onset. Here we instead applied the analysis on all trials irrespective of coherence. The CorCE results and preference for standard over delayed accumulation did not differ when using all the coherences. We quantified the shaded error bars using a bootstrap method with 100 resamples.

#### Population decoding

Data were preprocessed by calculating spike counts in 100 ms time windows, stepping every 20 ms, through the first 600 ms after motion onset, and again separately for spikes aligned to saccade onset (from 400 ms before to 200 ms after). We included only trials with RT > 400 ms to minimize edge effects that may obscure single-trial dynamics. We used two L1-regularized logistic decoders, one for choice and one for wager:

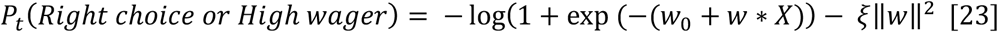

where *w* represents a vector of weights at time t (vector length matches number of units), *w*_*0*_ is a bias term, and *X* is the spike count vector for each unit in the session. The best hyperparameter ξ was found using a 50-value grid search between [0 1000], using 5-fold cross validation. The dataset was divided into a training set (90%) and test set (10%), the latter of which was used to calculate the prediction accuracy (Fig. 5b). If *P*_*t*_ was greater than 0.5 this would indicate that the decoder predicted either a right choice (for the choice decoder) or high bet (for the wager decoder). Values below 0.5 would indicate either a left choice or low bet. The performance (accuracy) was defined based on the monkey’s choice and wager at the end of the trial. We define a ‘neural decision variable’ (aka model DV; Kiani et al., 2014; Peixoto et al., 2021) by computing the log odds of a particular choice (e.g., 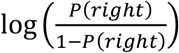) for rightward choices) given the population spike counts up to time *t* on a given trial (correct trials only). Importantly, for Figure 5a and 5c we used the log odds irrespective of choice, therefore the results combine 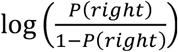 for right choices and 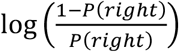 for left choices, to increase statistical power. For Figure 5c the log odds for wager are also combined in the following manner: 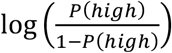 for high wagers and 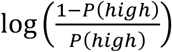 for low wagers.

To test whether the neural DV for the choice decoder showed a linear increase that was significantly dependent on motion strength, we fit a linear regression:

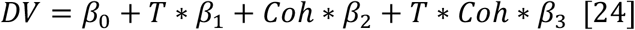

where *β*_0_ is a bias term, *T* is time (20 ms time bins for 200-600 ms after motion onset), *Coh* is motion coherence level, and β_*1*,*2*,*3*_ are the weights accompanying the predictor variables. If *β*_*3*_ was significantly different from zero *(P < 0*.*0*5), then the modulation of log odds by motion strength was deemed significant. To compute this linear regression we only used the mean *DV* shown in Figure 5a, excluding data cut off at the mean RT for each individual coherence.

As mentioned in the Results, both decoders provide weights for each neuron through time. Therefore, to test whether our sample of neurons comprised a single population that contributes approximately equally to choice and wager, we calculated the correlation between the weight magnitudes (irrespective of sign) as well as the distribution of the difference in weight magnitude. The weights were preprocessed by taking their absolute value because of our interest in magnitude and not direction. To calculate the correlation, we compute the Pearson’s correlation between absolute weights for all our neurons at each time point. We collapsed over time by averaging over the first 600 ms after motion onset and last 400 ms before saccade onset (Extended Fig. 5a, red line). We computed significance by randomly permutating the choice and wager weights 1000 times (Extended Fig. 5a, blue distribution). Additionally, the choice and wager decoder absolute weights were also subtracted from one another to create a distribution which informs whether there is a single population equally contributing to both choice and wager. Significance was evaluated using the Hartigan’s dip test, which tests whether a distribution is unimodal or bimodal.

To determine the trial-by-trial relationship between the choice and wager decoder we first tested whether the wager decoder was predictive of the DV (log odds). Trials were categorized as decoded-low or decoded-high confidence by calculating the mean *P*(*High*) for the wager decoder from 200 ms before saccade initiation until 100 ms after. Values above (below) .5 indicated a decoded-high (decoded-low) wager. We strictly looked at trials with 0% coherence to try and remove any effects of coherence. Results for Figure 6a were calculated by then averaging the DV on these decoded-high and decoded-low trials. Significance bars were calculated using a one-tailed t-test with Šidák’s correction. Importantly, differences in peak DV around saccade onset (Fig. 6a) highlight changes in the ratio 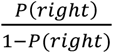.

To examine the relationship between the updating of choice and confidence at a finer time scale, we related the log odds of the choice decoder to the *P*(*High*) from the wager decoder on each individual trial. As shown in Figure 6a, there is a link between the model DV and *P*(*High*) that can be exploited to understand whether the temporal updating is in sync or if there is some lag. Figure 6e-g was generated using the 3 separate 400-ms time windows described in the legend. The independent variable, which is *P(right)* from the choice decoder, was corrected so that values near 0.5 (chance level) were close to zero and values moving away from 0.5 in either direction (better predicting left or right choice) became closer to one. This was done using the following equation:

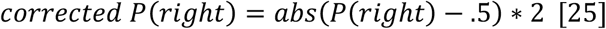

In essence, this transformation changes the property of choice decoding strength so that it is monotonic, making it possible for a linear relationship to exist between corrected *P(Right)* and *P*(*High*). To capture this relationship and its dependency on time lag, we applied a linear regression, using the corrected *P(Right)* at time *t*, to the dependent variable *P*(*High*) at time *t +* Δ*t*, where Δ*t* ranged from +/-200 ms. To quantify how informative corrected *P(Right)* is of *P*(*High*) we used an R^2^ measurement at each time lag. Figure 6e-g displays a corrected R^2^ which is simply the R^2^ value after subtracting out the average R^2^ values when the time series of the decoders for each trial were randomly permuted.

## Extended data figures for Vivar-Lazo & Fetsch

**Extended Data Fig. 1:**
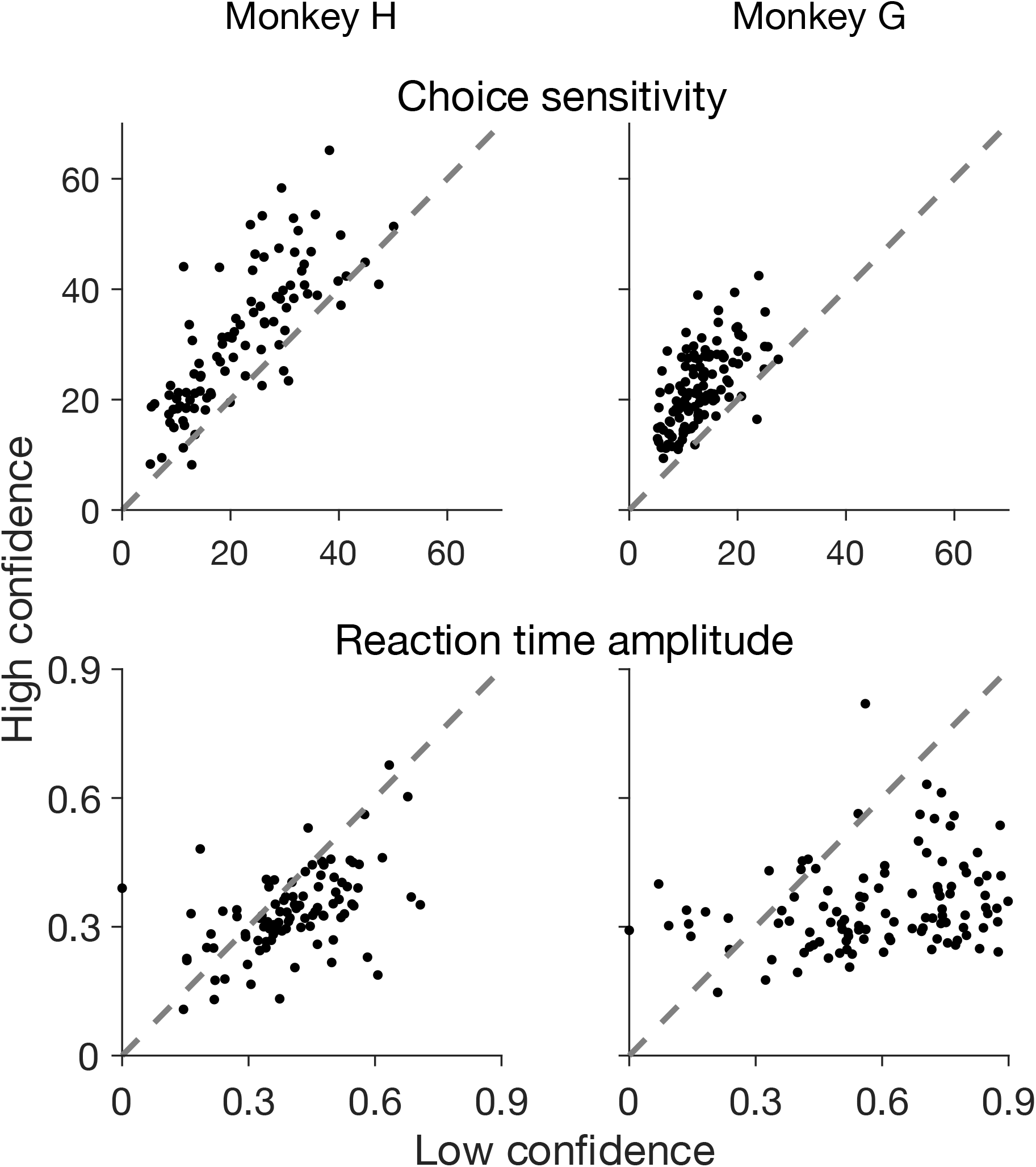
Session-by-session sensitivity and amplitude parameters from fitting logistic and Gaussian functions to choice and RT, respectively. The vast majority of individual sessions showed greater sensitivity (accuracy) and faster RT amplitude when the monkey bet high vs. low.

**Extended Data Fig. 2:**
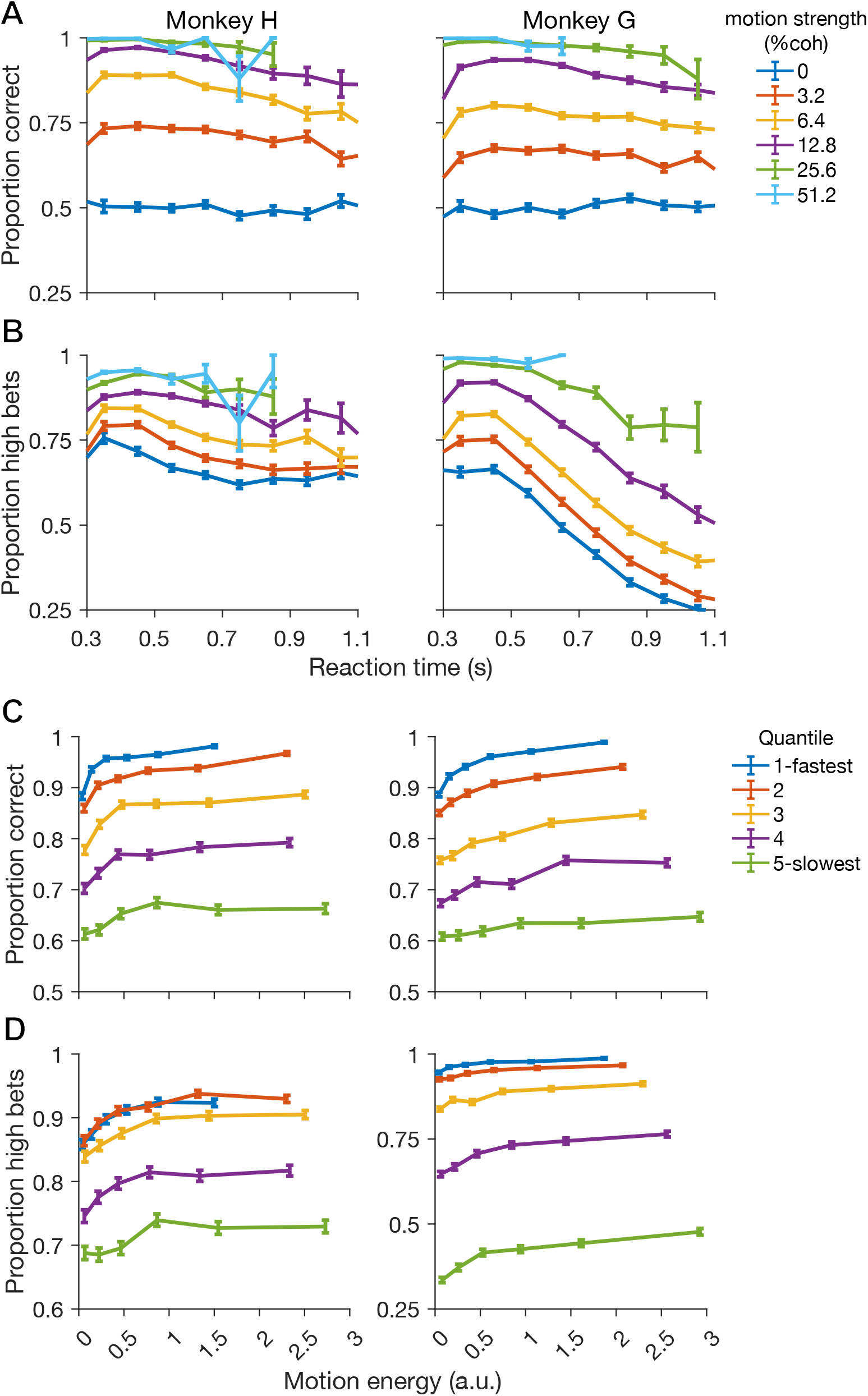
(A) Accuracy as a function of RT quantile, split by motion strength (% coh). Accuracy tended to decrease as a function of RT (*p* < .0085 for every coherence except 0% in both monkeys and 51.2% in monkey G). (B) Proportion of trials with a high bet as function of RT. Colors same as A. (C) Accuracy as a function of motion energy. Colors represents five different RT quantiles. Significant increases in accuracy were observed across all RT quantiles for both monkeys (p<0.01). (D) Proportion of trials with a high bet as a function of motion energy. Colors same as C.

**Extended Data Fig. 3:**
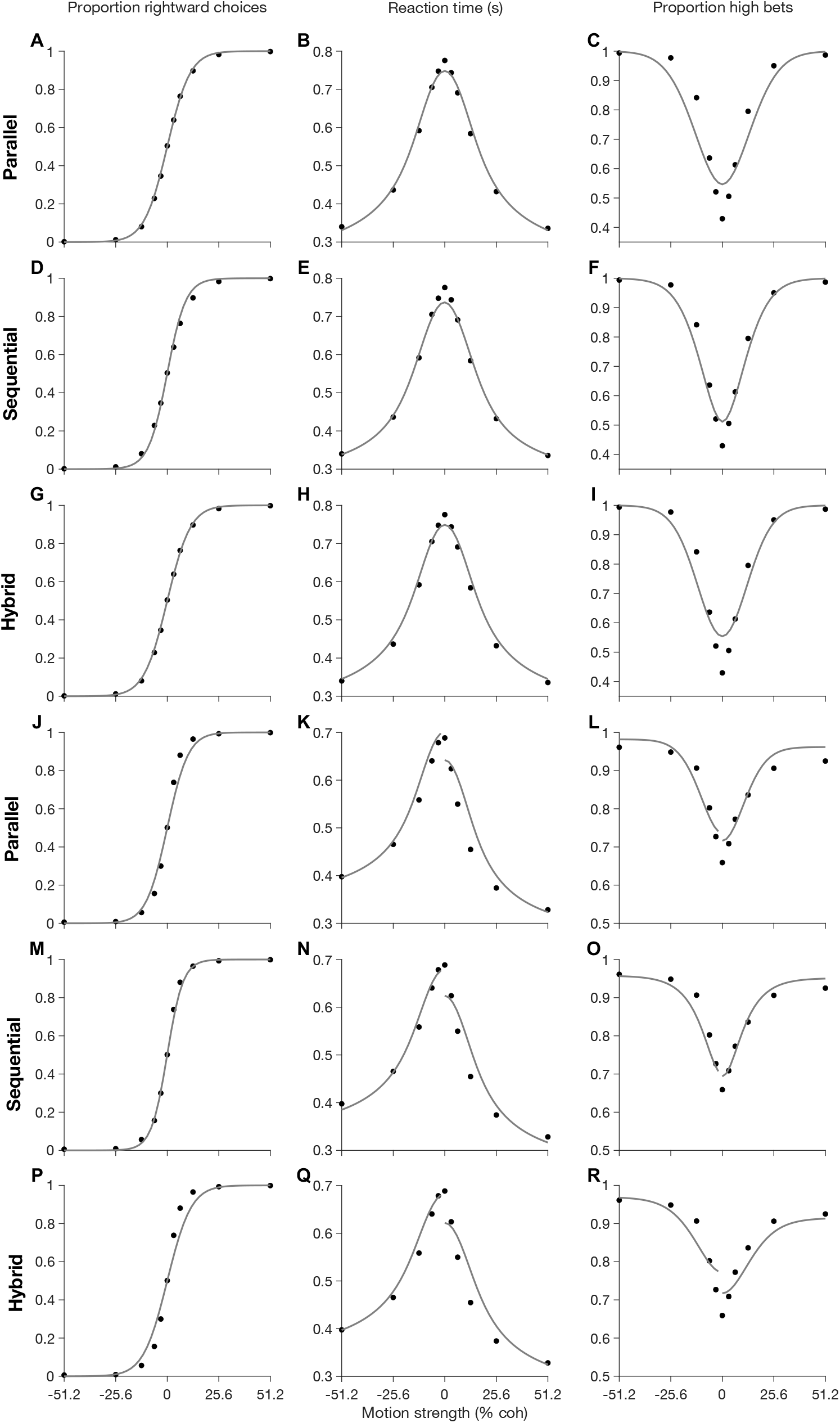
Serial, parallel, and hybrid model fits on unconditioned data. (A, B, C) Parallel model fitted to choice, RT, and wager as a function of motion strength (% coh). (D, E, F) Serial model fitted to choice, RT, and wager as a function of motion strength (% coh). (G, H, I) Hybrid model fitted to choice, RT, and wager as a function of motion strength (% coh). Figures A-I correspond to monkey G. (J, K, L) Parallel model fitted to choice, RT, and wager as a function of motion strength (% coh). (M, N, O) Serial model fitted to choice, RT, and wager as a function of motion strength (% coh). (P, Q, R) Hybrid model fitted to choice, RT, and wager as a function of motion strength (% coh). Figures J-R correspond to monkey H.

**Extended Data Fig. 4:**
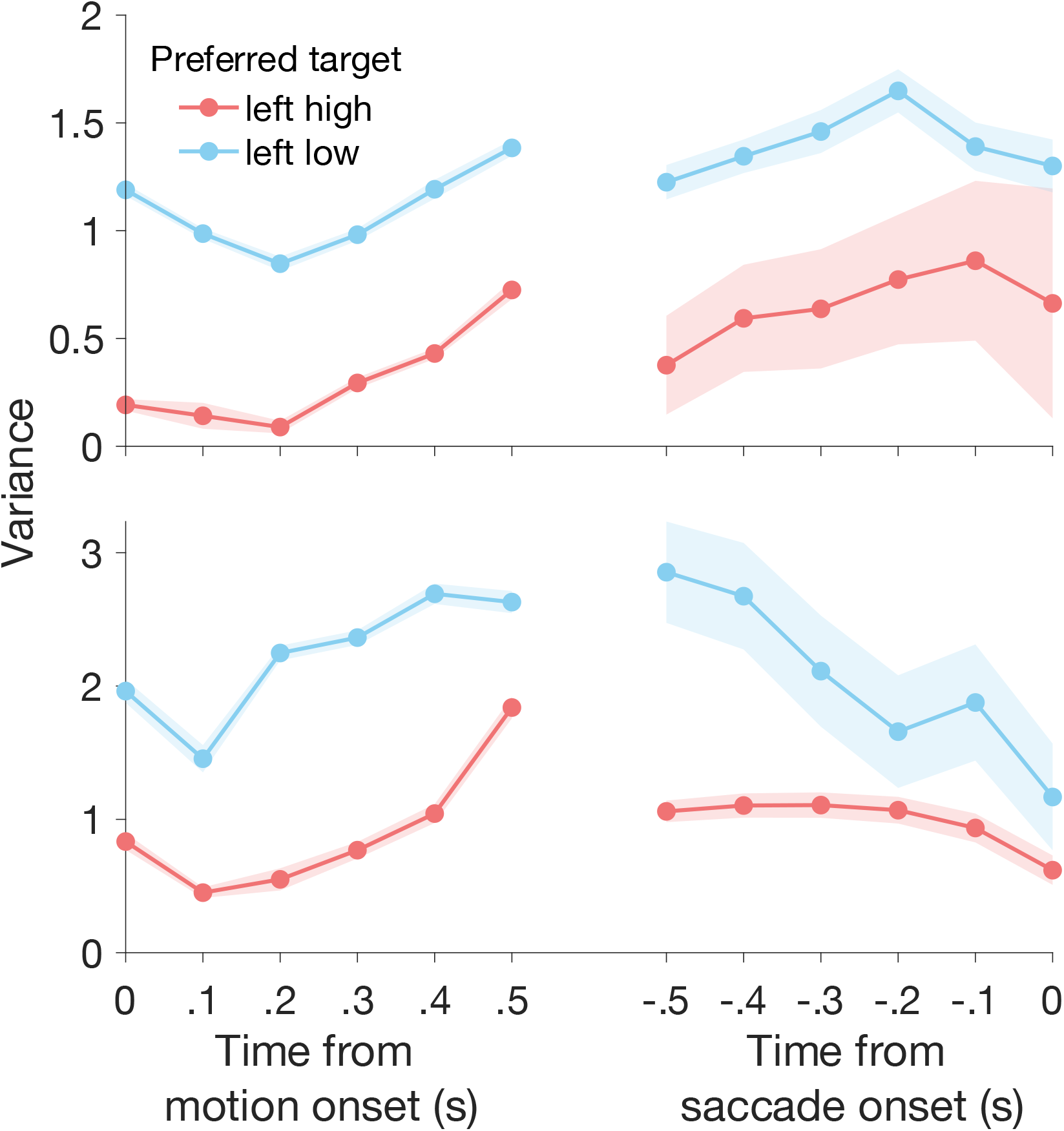
Variance of the conditional expectation (VarCE) estimated for the left-high and left-low populations of neurons. In both monkeys (H=top, G=bottom) VarCE begins to increase at approximately 0.2 s from motion onset, then decreases near saccade onset. Colors represent the two populations, and the shaded regions are standard errors calculated using a bootstrap.

**Extended Data Fig. 5:**
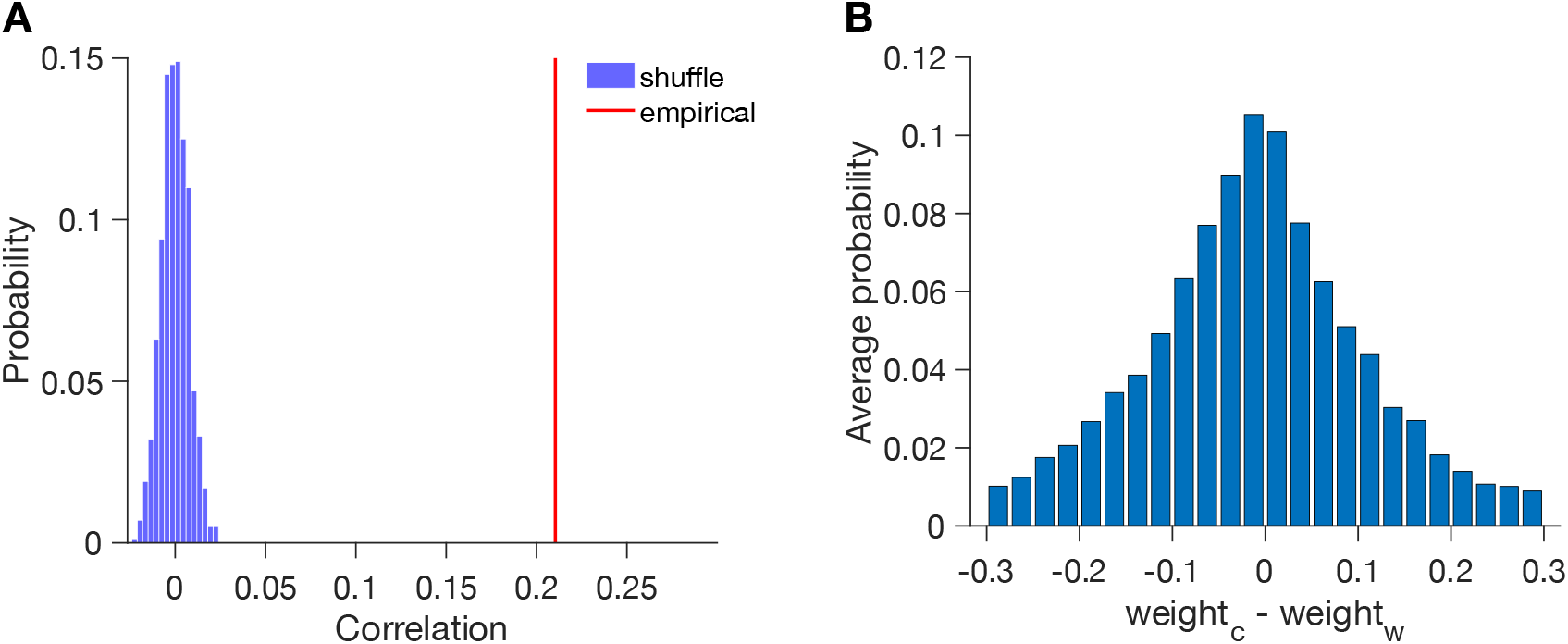
Correlation between population decoder weights. (A) Correlation between the magnitude of choice and wager decoder weights (red line) compared to the value expected by chance (shuffled data). (B) Histogram of the difference between choice and wager decoder weights. D and E show pooled data from both monkeys.

